# Multiplex RNA single molecule FISH of inducible mRNAs in single yeast cells

**DOI:** 10.1101/631622

**Authors:** Guoliang Li, Gregor Neuert

## Abstract

Transcript levels powerfully influence cell behavior and phenotype and are carefully regulated at several steps. Recently developed single cell approaches such as RNA single molecule fluorescence in-situ hybridization (smFISH) have produced advances in our understanding of how these steps work within the cell. In comparison to single-cell sequencing, smFISH provides more accurate quantification of RNA levels. Additionally, transcript subcellular localization is directly visualized, enabling the analysis of transcription (initiation and elongation), RNA export and degradation. As part of our efforts to investigate how this type of analysis can generate improved models of gene expression, we used smFISH to quantify the kinetic expression of *STL1* and *CTT1* mRNAs in single *Saccharomyces cerevisiae* cells upon 0.2 and 0.4M NaCl osmotic stress. In this Data Descriptor, we outline our procedure along with our data in the form of raw images and processed mRNA counts. We discuss how these data can be used to develop single cell modelling approaches, to study fundamental processes in transcription regulation and develop single cell image processing approaches.

## Background & Summary

Transcript levels are regulated by essential biological processes^1,2^. Traditionally, investigations of how these processes are regulated within the cell have relied on cell populated based approaches^3-5^. Such studies have uncovered important insights into how transcript levels are regulated. However, studies performed using recently developed single cell-based approaches have demonstrated that the population average reported from such experiments can unintentionally equate essentially distinct expression patterns^6-20^. For example, two cell populations, one wherein a few cells express high levels of a particular RNA and one wherein many cells express low levels of a particular RNA, would appear to have comparable RNA expression levels in population-based measurements. Additionally, cell populated methods are also unable to address whether two different mRNA species are expressed in the same cell or in different cells which has hampered the identification of distinct cell types. This transcript variability is problematic because any selection on these cells based on RNA expression levels could have a vastly different impact on these two cell populations in terms of cell vitality and/or the number of cells that survive.

In comparison to population measurements, single-cell/single-molecule imaging experiments such as RNA single molecule fluorescence in situ hybridization (smFISH) can resolve the spatial and temporal distribution of individual RNA molecules with high resolution (Figure 1). This enables the correlation of the expression of different RNA species with each other and with cellular phenotypes. Additionally, we and others have recently shown how RNA smFISH data can lead to better more predictive models of gene expression due to: 1) the fact that they contain information on more gene regulatory processes, including transcription (initiation, elongation), RNA export, and RNA degradation, and 2) they enable direct measurement of the distribution of RNA molecules which are essential for single cell modeling^20-27^.

**Figure 1.**
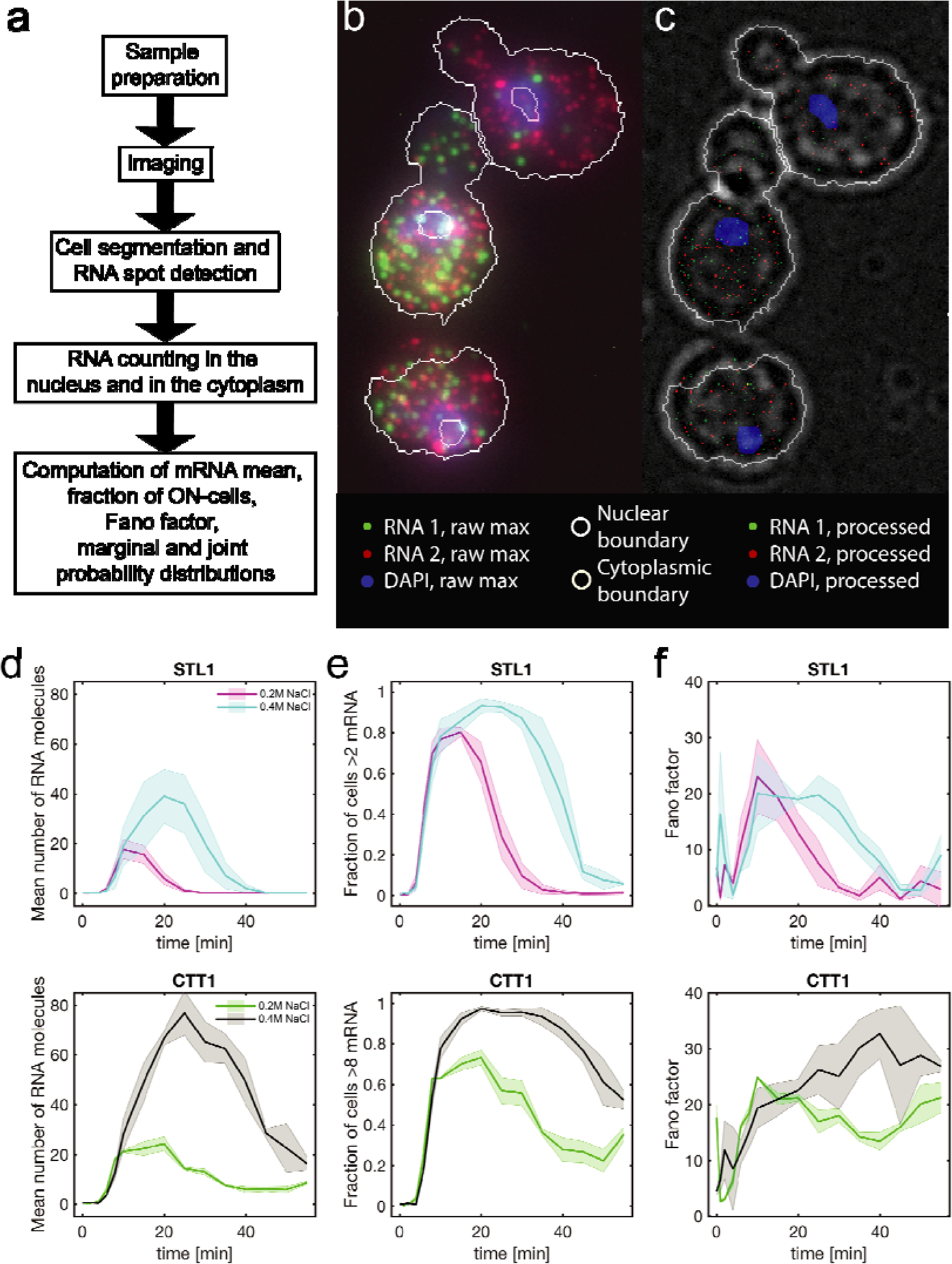
Generation and analysis of single molecule RNA fluorescent in-situ hybridization data sets. (a) Experimental and computational workflow. (b) Example image and (c) its analysis with segmented nuclear (grey) and cytoplasmic (grey) boundary, two RNA species (red and green spots) and DAPI stained nucleus (blue). Scale bar 5 µm. (d) Mean number of mRNA molecules, (e) fraction of ON-cells, and (f) Fano-factor for *STL1* and *CTT1* at 0.2M and 0.4M NaCl as a function of time. The solid lines and shaded areas represent the mean and standard deviations respectively from two biological replica experiments.

Unfortunately, very few RNA smFISH datasets are publicly available, and available datasets are limited in terms of the number or RNA species, time points, and cells. This limits the ability of the research community to develop improved RNA smFISH analysis approaches and discover new insights into the important biological processes regulating transcription levels.

To address this limitation, we present our information rich RNA smFISH data set in this Data Descriptor which is an extension to our previous publication^26,30^. The experimental and computational workflow consist of(1) sample preparation, (2) imaging cells, nucleus and RNA species, (3) nuclear and cytoplasmic segmentation, (4) RNA spot detection, (5) RNA counting and (6) computation of various sample statistics (Figure 1a). This data set reports the imaging and quantification of two mRNAs (*STL1* and *CTT1*) in 65,000 individual model yeast *Saccharomyces cerevisiae* cells imaged in 3D (Figures 1-3). The expression of these mRNAs was induced using two different osmotic stresses (0.2M and 0.4M NaCl) and monitored over sixteen time points in biological duplicate or triplicate^26, 28^.

Because this data set contains both 3D spatial and temporal information on the expression of these RNAs, it can be used to simultaneously investigate many different important processes regulating transcription levels as they occur in the cell. Such analyses are not possible in cell population-based or single cell sequencing experiments^1-5,27^. Here we present RNA expression as population mean, fraction of cells above basal mRNA expression (ON-cells), the variance normalized by the expression mean (Fano factor) (Figure 1), marginal probability of nuclear and cytoplasmic mRNA (Figure 2), and the joint probability of nuclear and cytoplasmic RNA expression (Figure 3).

**Figure 2.**
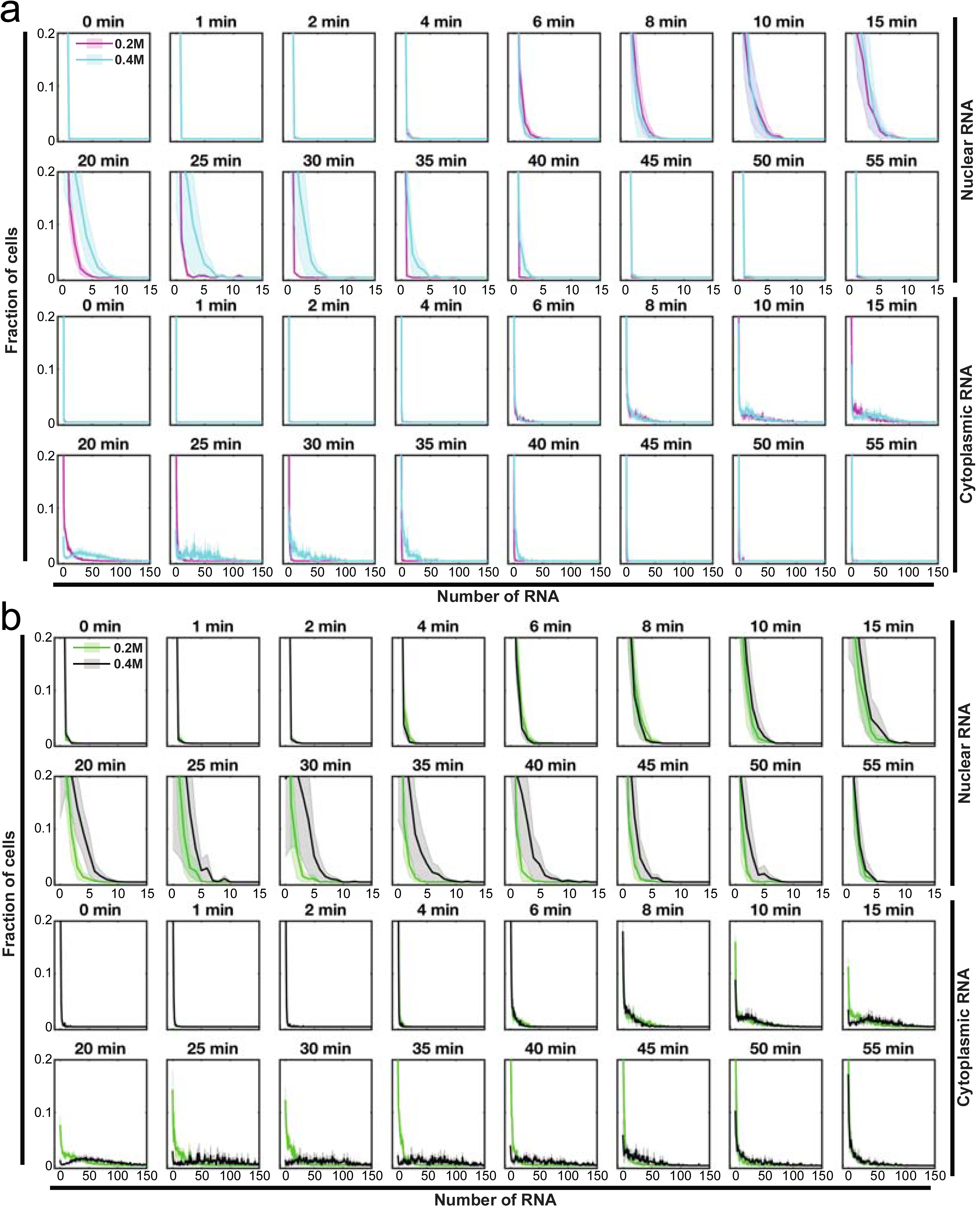
Single cell marginal mRNA distributions as a function of time. (a) *STL1* and (b) *CTT1* mRNA expression in the nucleus (top) and cytoplasm (bottom) at 0.2M and 0.4M NaCl at sixteen different time points. The solid line and shaded area represent the mean and standard deviations from two (0.2M NaCl) or three (0.4M NaCl) biological replica experiments.

**Figure 3.**
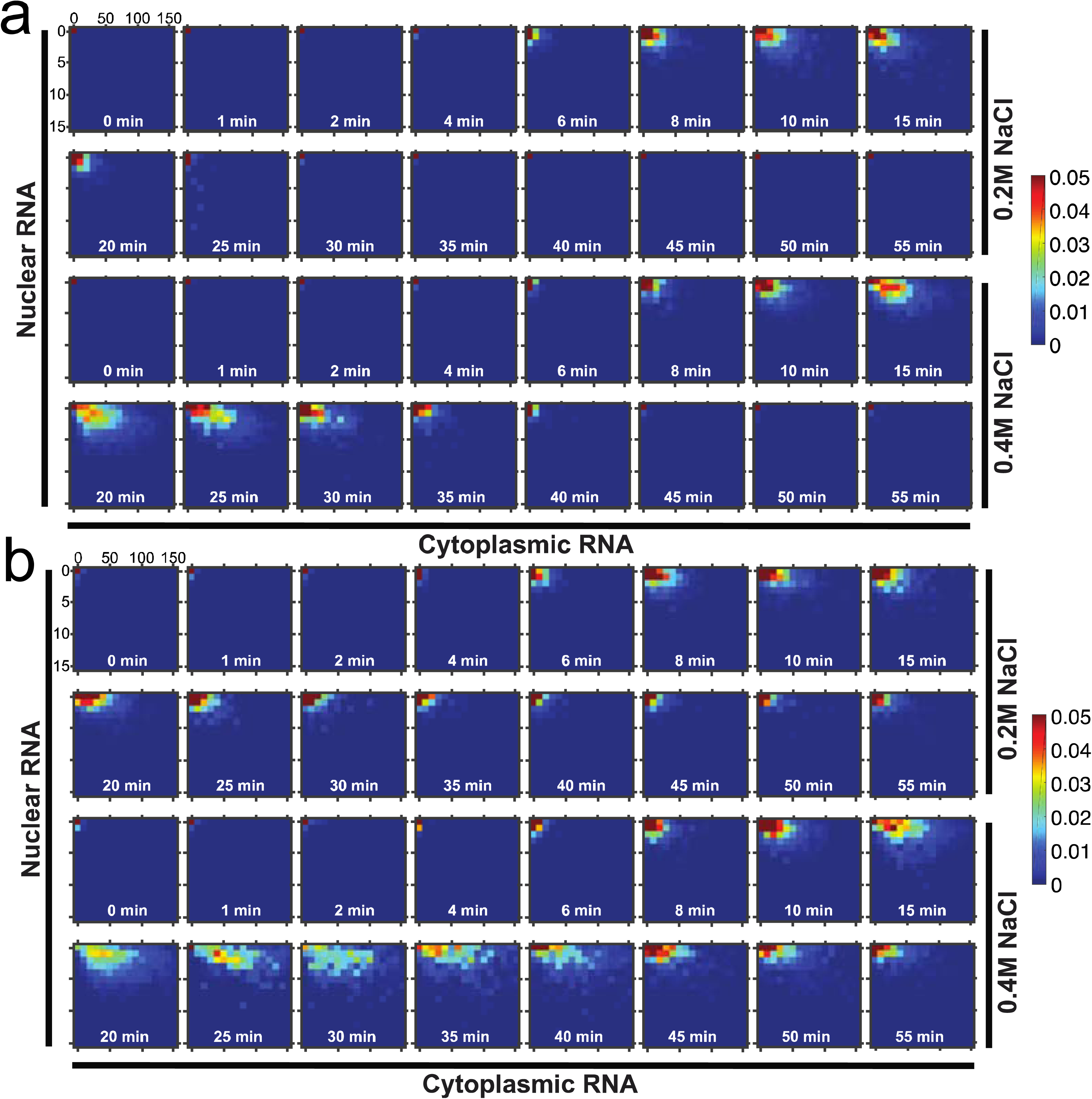
Single cell joint probability distribution of nuclear and cytoplasmic mRNA as a function of time. (a) *STL1* and (b) *CTT1* mRNA joint probability distribution of the nucleus (vertical axis) and cytoplasm (horizontal axis) at 0.2M (top) and 0.4M (bottom) NaCl at sixteen different time points. The mean from two biological replica is plotted. Color encodes probability (0 – 0.05 = 0-5%).

The reuse potential of this data set lies in three independent areas of research. The first area is in gene expression modelling. Standard approaches to modelling do not always work well on single cell data sets. This data set could be particularly helpful for theoretical labs working to develop improved modelling approaches and which may not have the resources or expertise to generate their own similar data sets. The second area is in image processing. The limit on the amount and quality of data that can be extracted from data sets such as these is set by image processing. Thus, improved image processing approaches are needed to maximize the amount of data extracted from one data set. Finally, we expect that such a dataset could be re-used by labs interested in studying transcription, RNA export, and RNA degradation as these processes have not yet been thoroughly investigated using single cell-based approaches.

## Methods

These methods are expanded versions of descriptions in our related work^26^.

### Yeast strain and growth condition

*Saccharomyces cerevisiae* strain *BY4741* (*MATa his3*Δ*1 leu2*Δ*0 met15*Δ*0 ura3*Δ*0*) was used for RNA smFISH experiments. Three days before the experiment, yeast cells were streaked out on a complete synthetic media plate (CSM, Formedia, UK) from a glycerol stock stored at −80 C. The day before the experiment, a single colony from the CSM plate was inoculated in 5 ml CSM medium (pre-culture). After 6-12h, the optical density (OD) of the pre-culture was measured and the cells were diluted into new CSM medium to reach an OD of 0.5 the next morning.

### RNA single molecule Fluorescence In-Situ Hybridization (smFISH) Probe Construction

Probes used to detect discrete *STL1* and *CTT1* mRNAs consist of a set of 48 DNA oligonucleotides, each with a length of 20nt (see Table 1 for DNA sequences). The 3’-end of each probe was modified with an amine group and coupled to either tetramethylrhodamine (TMR) (Invitrogen) (*STL1*) or Cy5 (GE Amersham) (*CTT1*). Coupled probes were ethanol precipitated and purified on an HPLC column to isolate oligonucleotides displaying the highest degree of coupling of the fluorophore to the amine groups.

**Table 1.**
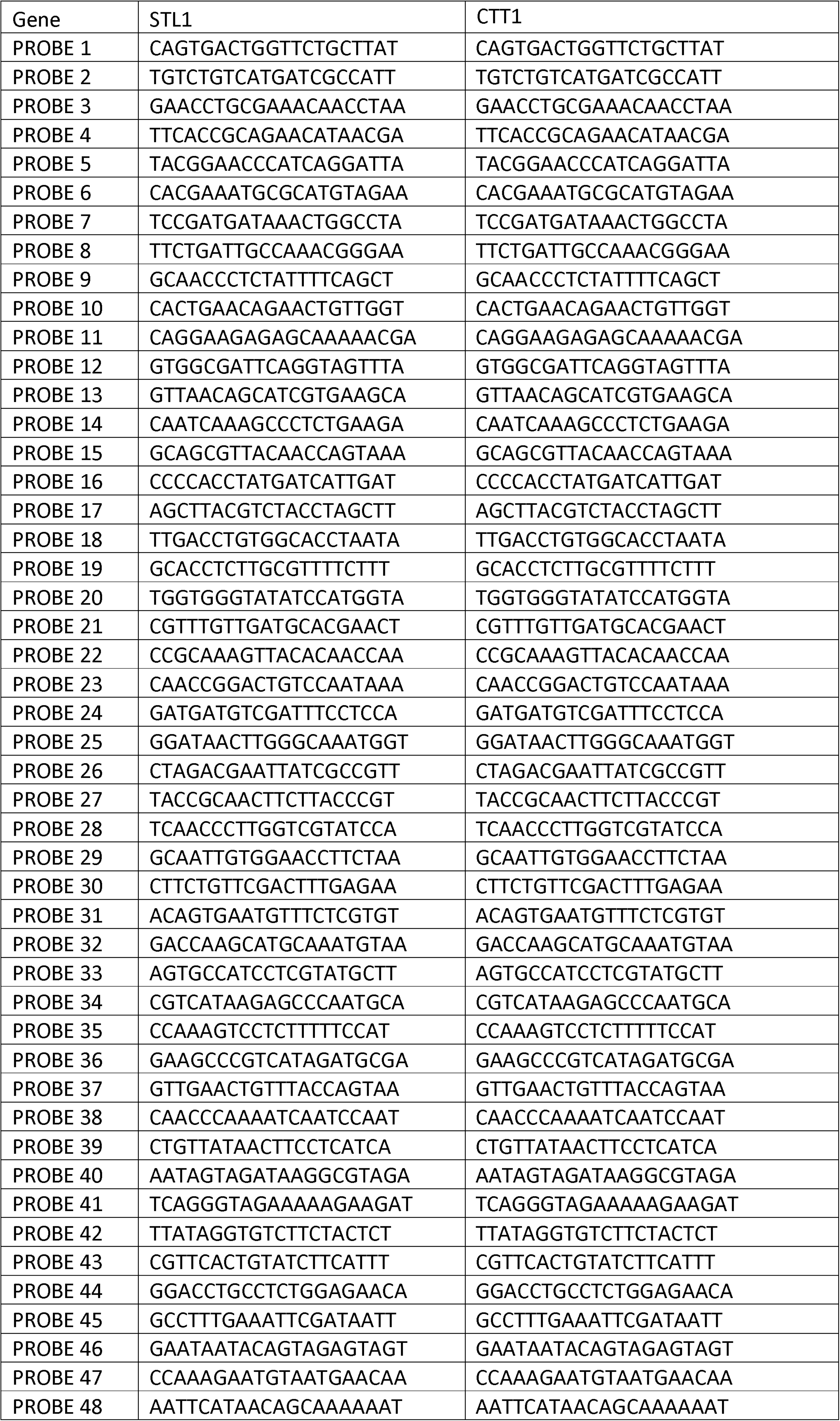
RNA-FISH probes.

### Sample preparation for smRNA-FISH

Yeast cells in log-phase growth (OD = 0.5) were concentrated 10 times (OD = 5) by a glass filter system with a 0.45 μm filter (Millipore). Cells were exposed to a final osmolyte concentration of either 0.2 M or 0.4 M NaCl, and fixed in 4 % formaldehyde at time points 0, 1, 2, 4, 6, 8, 10, 15, 20, 25, 30, 35, 40, 45, 50, and 55 minutes. At each time point, cells (5 ml) in the corresponding beaker were poured into a 15 ml falcon tube containing 0.55ml of 37% formaldehyde, resulting in cell fixation. Cells were fixed at +20C for 30 minutes, then transferred to +4C and fixed overnight on a shaker. After fixation, cells were centrifuged at 500×g for 5 minutes. The liquid phase was discarded, the cell pellet was resuspended in 5 ml ice-cold Buffer B (1.2 M sorbitol, 0.1 M potassium phosphate dibasic, pH 7.5), and cells were centrifuged again. After discarding the liquid phase, yeast cells were resuspended in 1 ml Buffer B, and transferred to 1.5 ml centrifugation tubes. Cells were then centrifuged again at 500×g for 5 minutes. The supernatant was removed and the pellet was resuspended in 0.5 ml Spheroplasting Buffer (Buffer B, 0.2 % β-mercaptoethanol, 10 mM Vanadyl-ribonucleoside complex). The OD of each sample was measured and samples were adjusted such that each contained the same number of cells. 10 μl of 2.5 mg/ml Zymolyase (US Biological) was added to each sample on a +4C block. Cells were incubated on a rotor at +30C and cell wall digestion was monitored by microscopy. Zymolyase digestion was terminated when 90% of the cells visualized under the microscope were black instead of opaque (indicating cell wall digestion) by incubation on a +4C block. Cells were centrifuged for 5 minutes, then the cell pellet was resuspended with 1 ml ice-cold Buffer B and spun down for 5 minutes at 500×g. After discarding the liquid phase, the pellet was gently re-suspended with 1 ml of 70 % ethanol and kept at +4C for at least 12 hours at +4C. Cells were incubated with a 1:1000 dilution (chosen for optimal ratio of RNA spot intensity to cell background as determined by independent optimization experiments) of RNA smFISH probes (stock concentration: STL1-TMR probes: 242pmol/µl; CTT1-Cy5 probes: 90 pmol/µl) in hybridization buffer (10% dextran, 10 mM vandyl-ribonucleoside complex, 0.02% RNAse-free BSA, 1mg /ml E.coli tRNA, 2x SSC, 10% formamide) to label *STL1* and *CTT1* mRNAs.

### Microscopy setup for single molecule RNA FISH imaging

Cells were imaged with a Nikon Ti Eclipse epifluorescent microscope equipped with perfect focus (Nikon), a 100× VC DIC lens (Nikon), fluorescent filters for DAPI, TMR and CY5 (Semrock), an X-cite 120 fluorescent light source (Excelitas), and an Orca Flash 4v2 CMOS camera (Hamamatsu). The microscope was controlled by the Micro-Manager software program^29^.

### Image acquisition for single molecule RNA FISH imaging

For single molecule RNA-FISH microscopy, z-stacks of images from fixed yeast cells for widefield (20ms exposure), DAPI (20ms exposure), TMR (1s exposure) and CY5 (1s exposure) were taken with each image in the z-stack separated by 200 nm. For each sample, multiple positions on the slide were imaged to ensure large numbers of cells.

### Image analysis for single-molecule RNA-FISH

RNA smFISH images generated by microscopy were analyzed to segment cells and define the nuclear and cytoplasmic compartment of each cell^26,32^. For each DAPI image stack, a maximum intensity projection was generated. A DAPI intensity threshold was picked by eye to identify the maximum number of nuclei in the image. Based on this threshold, the image was then converted into a binary image. Connected regions that were too big or too small were removed. Connected regions in the image were then labeled by individual numbers resulting in individual numbered nuclei. Next, for each connected region an individual DAPI threshold was computed as 50% of the difference between the maximum DAPI signal minus the background DAPI signal. Through this iterative and cell specific thresholding, cell-to-cell differences in nuclei DNA concentration and DAPI staining were considered. For each cell, the entire DAPI image stack was converted into a binary image stack using the cell-specific DAPI thresholds, resulting in a 3D nucleus and cytoplasm where thresholds have been considered. After the nucleus had been segmented, the last 5 images of the widefield image stack were subjected to maximum intensity projection to generate the cell outlines. A background image was generated by running a disk smoothing filter over this image. The background image was subtracted from the maximum projected image to enhance contrast. This image was then combined with the thresholded DAPI image using morphological reconstruction and cells were segmented with a watershed algorithm. After image segmentation, elements that were not properly segmented and resulted in artificially small or large segmented elements were removed. Cells too close to the image borders were also removed.

To identify RNA spots, fluorescence thresholds for RNA-FISH images were determined for each dye based on a single image plane, for each position imaged and for each sample. For a given dye, the mean threshold value was calculated based on the thresholds identified from each image in the data set. This threshold was reproducible from image to image and from person to person. After the threshold had been determined for each dye, each image in the stack was subjected to a Gaussian filter to remove noise, and then filtered with a Laplacian of a Gaussian filter to detect spots in the image. These filtered images were then converted into binary images using the previously determined threshold. For each pixilated signal, the regional maximum was determined, which identifies the xyz-position of the RNA spot within each 3D image stack. This process was repeated for each of the TMR and CY5 image stacks. The number of RNA spots for each cell in the cytoplasm or the nucleus was determined by applying the mask of segmented nucleus and cytoplasm on the filtered RNA-FISH image stacks in 3D. The numbers of RNA molecules in the nucleus and cytoplasm were counted for each individual cell. The RNA-FISH data set consists of a total of 65454 single cells (25511 at 0.2M NaCl and 39943 at 0.4M NaCl).

### RNA-FISH data analysis

For each time point, a distribution of mRNA molecules in single cells was determined as the marginal distribution of total *STL1* and *CTT1* mRNA or as the joint probability distribution of nuclear and cytoplasmic RNA. The distributions were also summarized in a binary data set of ON and OFF cells. ON-fraction are considered cells that turn on RNA transcription after osmotic stress. RNA-expression prior to osmotic stress is considered basal RNA transcription of the gene and is considered an OFF-cell. For the quantification of ON-fraction, the basal expression defined as 95% of the cumulative *STL1* and *CTT1* RNA distribution was chosen resulting in ON-cells with more than two (*STL1*) or more than eight (*CTT1*) mRNA molecules. The population average was computed as the mean marginal cytoplasmic or nuclear RNA for each mRNA species.

### Code availability

The computer codes used to generate spatiotemporal RNA expression data sets reported in this study and for automated cell segmentation and RNA spot detection were developed at Vanderbilt University and are intended solely for scientific research. The developers of the code are willing to apply the algorithms to data sets provided by the research community through academic collaboration. Interested readers should contact the corresponding author for further details.

### Data Records

The datasets are available from the Image Data Resource (IDR)^30^, under the following DOI: https://doi.org/10.17867/10000118.

Data also have been deposited to BioStudies with the dataset identifier S-BIAD1, accessible through the URL: https://www.ebi.ac.uk/biostudies/studies/S-BIAD1.

Data also have been deposited to a public dropbox folder (https://www.dropbox.com/sh/4ydp2dnnu8kvdrm/AAA3eDV5CJOllWJprvwEjGy8a?dl=0) The data sets consist of raw image stacks, analysed images and RNA spot counts in each cell.

Statistical analysis using a ks-test was performed on marginal distributions and the test statistics are available within Figshare^31^

### Raw datasets: widefield and fluorescent microscopy image stack

Each experimental condition, time point, and field of view is stored as an image stack consisting of 25 or 26 images of each *STL1* mRNA labelled with TMR coupled oligonucleotides, *CTT1* labelled with CY5 coupled oligonucleotides (see Table 1 for the DNA sequences), the DAPI stained DNA, and widefield images of the cells. Each stack consists of 100 or 104 images and is stored as a TIFF file. In total, 4-8 fields of view have been imagined at 16 different time points, for two experimental conditions each in biological duplicates or triplicates (331 image stacks = 277.44GB) and can be accessed on-line (Table 2).

**Table 2:** Experimental data sets and conditions.

| Experimental condition | Time points [minutes] | Stacks of images (size) | Link to data sets |
| --- | --- | --- | --- |
| 0.2M NaCl, step, replica 1 (Exp1_rep1) | 0, 1, 2, 4, 6, 8, 10, 15, 20, 25, 30, 35, 40, 45, 50, 55 | 65 (55.46 GB) | https://idr.openmicroscopy.org/webclient/?show=project-503<br>https://www.ebi.ac.uk/biostudies/studies/S-BIAD1 |
| 0.2M NaCl, step, replica 2 (Exp1_rep2) | 0, 1, 2, 4, 6, 8, 10, 15, 20, 25, 30, 35, 40, 45, 50, 55 | 65(53.71 GB) | https://idr.openmicroscopy.org/webclient/?show=project-503<br>https://www.ebi.ac.uk/biostudies/studies/S-BIAD1 |
| 0.4M NaCl, step, replica 1 (Exp2_rep1) | 0, 1, 2, 4, 6, 8, 10, 15, 20, 25, 30, 35, 40, 45, 50, 55 | 72 (59.66 GB) | https://idr.openmicroscopy.org/webclient/?show=project-503<br>https://www.ebi.ac.uk/biostudies/studies/S-BIAD1 |
| 0.4M NaCl, step, replica 1 (Exp2_rep2) | 0, 1, 2, 4, 6, 8, 10, 15, 20, 25, 30, 35, 40, 45, 50, 55 | 68 (56.23 GB) | https://idr.openmicroscopy.org/webclient/?show=project-503<br>https://www.ebi.ac.uk/biostudies/studies/S-BIAD1 |
| 0.4M NaCl, step, replica 3 (Exp2_rep3) | 0, 2, 4, 6, 8, 10, 15, 20, 25, 30, 35, 40, 45, 50, 55 | 61 (52.38 GB) | https://idr.openmicroscopy.org/webclient/?show=project-503<br>https://www.ebi.ac.uk/biostudies/studies/S-BIAD1 |

### Analysed images

For each image stack the segmented and the processed RNA tif images are provided and they are described in Table 3. In addition, summary files of RNA counts per cell are available as CSV files^30^.

**Table 3:**
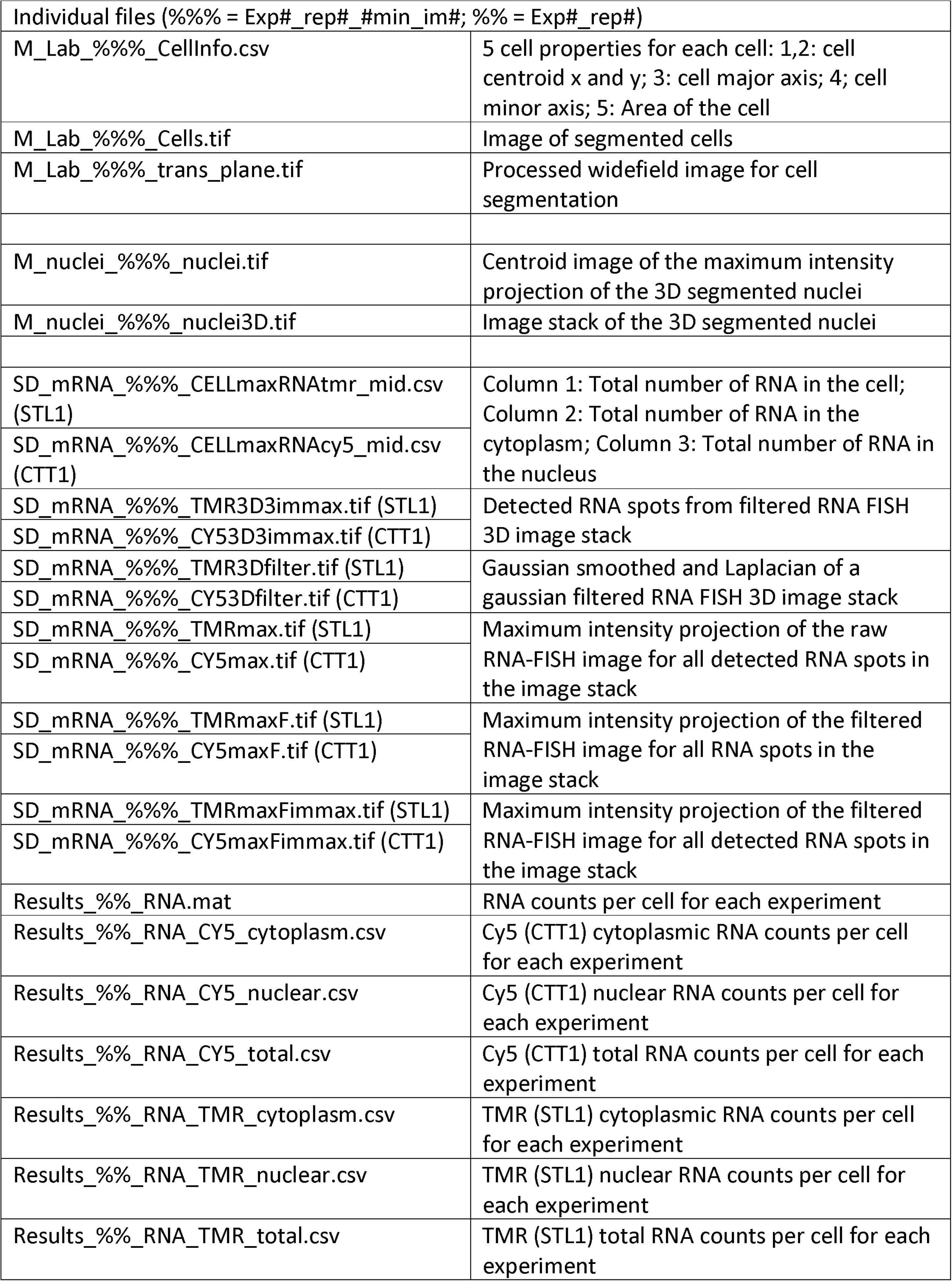
Content of the MATLAB and exported files (D = dimension of the array).

### RNA spot counts in each cell

After all the images for an experimental data set have been analysed, the analysed data from all image stacks are combined into a single file for each experiment. The content of this file is described in Table 3 and is available as MATLAB and CVS files. From these datasets, the RNA counts per cell are then used to compute the mean (Figure 1c), fraction of ON-cells (Figure 1d), Fano factor (Figure 1e), marginal (Figure 2) and joint probability distribution (Figure 3). Statistical analysis using a ks-test was performed on marginal distributions^31^.

### Technical Validation

#### Experimental validation

During sample preparation, the cell wall of the yeast cells needs to be digested. If the cell wall is digested correctly, cells turn from an opaque to a black color. Cells were monitored under the microscope every 10 minutes after addition of Zymolyase and when 90 % of the cells turned black, cells were transferred to the +4C block to stop Zymolyase activity.

Optimizing RNA-FISH probe concentration (dilutions: 1:500; 1:1000; 1:2000; 1:4000; 1:8000) and formamide concentration (5, 10, 15, 20, 25 %) in the hybridization buffer is an important step in validating the RNA-FISH methodology. We use hybridization conditions without probes as a control in these experiments to determine any background signal. At time t = 0 min, mRNA expression of *STL1* is very low as previously shown with RNA-FISH and qRT-PCR^15^.

#### Validation of the computational pipeline

Each data set contains 122 to 232 images per stack. For each image stack the initial threshold to identify the maximum number of DAPI stained nuclei was determined manually by comparing the raw image with the image of the identified nuclei. Similarly, the thresholds for RNA spot detection in the TMR and CY5 channel was determined manually for each image stack by comparing the raw image with the image of the identified TMR or CY5 spots and then the mean RNA-spot threshold was calculated.

### Usage Notes

#### Raw datasets: widefield and fluorescent microscopy image stack

These datasets could be re-used to develop new and/or improved image processing applications. Each stack contains all the images acquired at a specific field of view. The resolution of the images is 2048 pixels by 2048 pixels. The number of images per stack is 100 or 104 with 25 or 26 z-position’s. At each z-position, a TMR (image 1), a CY5 (image 2), a DAPI (image 3), and a widefield (image 4) were acquired.

#### Analysed images

These datasets could be used to compare different image processing applications during application development and for investigating processes that affect the spatial organization of mRNA in the cell.

#### RNA spot counts in each cell

The data sets for each experiment contain the number of mRNA molecules in each cell, the cytoplasm, or the nucleus for each of the mRNA species visualized. These data sets can be directly used to compute the mean, the ON-fraction or the Fano factor. The Fano factor is computed as the variance divided by the mean. In addition, these data sets can be used to compute marginal nuclear and cytoplasmic RNA distributions as well as the joint probability distribution of nuclear and cytoplasmic RNA for *STL1* and *CTT1*.

## Acknowledgements

GL and GN were funded by NIH DP2 GM11484901, NIH R01GM115892 and Vanderbilt Startup Funds. The authors thank Alexander Thiemicke, Benjamin Kesler, Hossein Jashnsaz, and Amanda Johnson for comments on the manuscript.

## Author contributions

GL generated the data and ran the image processing. GN wrote the data analysis pipeline, generated the figures and wrote the manuscript.

