## Supplemental Table 1-3 for "Multiplex RNA single molecule FISH of inducible mRNAs in single yeast cells"

**Table S1: p-value from ks-test comparing different osmotic stress condition.**

| Gene | STL1 | | | |
|---|---|---|---|---|
| Conbdition | 0.2M vs. 0.4M | | | |
| Nuc / Cyto | Nuc | | Cyto | |
| Replica | 1 | 2 | 1 | 2 |
| 0 min | 1.0E+00 | 1.0E+00 | 1.0E+00 | 1.0E+00 |
| 1 min | 1.0E+00 | 1.0E+00 | 1.0E+00 | 1.0E+00 |
| 2 min | 1.0E+00 | 1.0E+00 | 1.0E+00 | 1.0E+00 |
| 4 min | 1.0E+00 | 1.0E+00 | 9.7E-01 | 8.0E-01 |
| 6 min | 1.0E-02 | 1.8E-01 | 4.6E-03 | 7.0E-01 |
| 8 min | 1.7E-47 | 1.7E-02 | 1.9E-140 | 3.4E-03 |
| 10 min | 6.7E-02 | 5.9E-02 | 1.9E-07 | 1.0E-04 |
| 15 min | 2.9E-171 | 4.8E-01 | 0.0E+00 | 8.5E-01 |
| 20 min | 0.0E+00 | 5.9E-137 | 0.0E+00 | 1.3E-260 |
| 25 min | 0.0E+00 | 8.5E-134 | 0.0E+00 | 1.2E-266 |
| 30 min | 0.0E+00 | 3.3E-148 | 0.0E+00 | 7.0E-255 |
| 35 min | 0.0E+00 | 1.5E-128 | 0.0E+00 | 3.1E-275 |
| 40 min | 0.0E+00 | 3.2E-96 | 0.0E+00 | 2.5E-224 |
| 45 min | 0.0E+00 | 5.4E-85 | 0.0E+00 | 5.4E-165 |
| 50 min | 0.0E+00 | 6.4E-49 | 0.0E+00 | 1.4E-90 |
| 55 min | 0.0E+00 | 4.2E-49 | 0.0E+00 | 1.4E-90 |

| Gene | CTT1 | | | |
|---|---|---|---|---|
| Conbdition | 0.2M vs. 0.4M | | | |
| Nuc / Cyto | Nuc | | Cyto | |
| Replica | 1 | 2 | 1 | 2 |
| 0 min | 1.0E+00 | 1.0E+00 | 4.3E-02 | 8.3E-01 |
| 1 min | 1.0E+00 | 1.0E+00 | 6.8E-01 | 5.6E-03 |
| 2 min | 1.0E+00 | 1.0E+00 | 2.4E-02 | 1.1E-01 |
| 4 min | 1.0E+00 | 4.1E-03 | 2.5E-01 | 4.7E-03 |
| 6 min | 2.0E-02 | 7.6E-03 | 6.4E-02 | 4.2E-03 |
| 8 min | 1.5E-41 | 7.8E-05 | 1.5E-120 | 7.1E-05 |
| 10 min | 2.9E-03 | 3.4E-01 | 1.5E-10 | 2.5E-01 |
| 15 min | 3.0E-139 | 1.9E-01 | 0.0E+00 | 4.9E-01 |
| 20 min | 0.0E+00 | 1.0E-139 | 0.0E+00 | 4.6E-182 |
| 25 min | 2.7E-250 | 1.6E-140 | 0.0E+00 | 2.6E-192 |
| 30 min | 0.0E+00 | 7.4E-142 | 0.0E+00 | 3.8E-198 |
| 35 min | 0.0E+00 | 1.0E-187 | 0.0E+00 | 2.5E-259 |
| 40 min | 2.6E-302 | 1.6E-182 | 0.0E+00 | 4.2E-260 |
| 45 min | 6.1E-304 | 9.5E-163 | 0.0E+00 | 8.7E-233 |
| 50 min | 1.1E-295 | 1.4E-111 | 0.0E+00 | 7.6E-190 |
| 55 min | 5.4E-243 | 2.5E-77 | 0.0E+00 | 6.9E-127 |

**Table S2: p-value from ks-test comparing different replica experiments.**

| Gene | STL1 | | | |
|---|---|---|---|---|
| Condition | 0.2M | 0.4M | 0.2M | 0.4M |
| Compartment | Nucleus | | Cytoplasm | |
| Replica | 1 vs 2 | | | |
| 0 min | 1.0E+00 | 1.0E+00 | 1.0E+00 | 1.0E+00 |
| 1 min | 1.0E+00 | 1.0E+00 | 1.0E+00 | 1.0E+00 |
| 2 min | 1.0E+00 | 1.0E+00 | 1.0E+00 | 1.0E+00 |
| 4 min | 1.0E+00 | 1.0E+00 | 3.4E-01 | 1.0E+00 |
| 6 min | 1.0E+00 | 7.7E-01 | 9.4E-01 | 3.1E-07 |
| 8 min | 7.4E-20 | 9.7E-01 | 2.5E-67 | 5.3E-09 |
| 10 min | 1.1E-17 | 1.2E-30 | 2.9E-34 | 9.2E-56 |
| 15 min | 4.0E-16 | 3.1E-261 | 3.0E-45 | 0.0E+00 |
| 20 min | 7.1E-04 | 1.7E-55 | 1.2E-12 | 7.3E-46 |
| 25 min | 5.3E-01 | 1.0E-38 | 3.9E-03 | 7.8E-48 |
| 30 min | 9.8E-01 | 1.6E-30 | 2.0E-07 | 3.1E-32 |
| 35 min | 8.1E-01 | 4.6E-47 | 1.7E-01 | 2.7E-65 |
| 40 min | 8.4E-01 | 1.9E-44 | 6.6E-01 | 3.4E-62 |
| 45 min | 1.0E+00 | 2.6E-46 | 8.0E-03 | 3.4E-63 |
| 50 min | 1.0E+00 | 2.9E-79 | 3.7E-03 | 2.5E-98 |
| 55 min | 7.3E-01 | 1.5E-79 | 2.6E-09 | 3.6E-104 |

| Gene | CTT1 | | | |
|---|---|---|---|---|
| Conbdition | 0.2M | 0.4M | 0.2M | 0.4M |
| Compartment | Nucleus | | Cytoplasm | |
| Replica | 1 vs 2 | | | |
| 0 min | 1.0E+00 | 1.0E+00 | 1.0E+00 | 1.0E+00 |
| 1 min | 1.0E+00 | 1.0E+00 | 7.4E-02 | 7.1E-01 |
| 2 min | 1.0E+00 | 1.0E+00 | 6.4E-01 | 2.7E-02 |
| 4 min | 4.8E-02 | 1.0E+00 | 9.5E-01 | 9.8E-03 |
| 6 min | 5.0E-04 | 1.3E-02 | 1.1E-08 | 4.6E-03 |
| 8 min | 1.9E-03 | 1.9E-03 | 3.6E-39 | 8.9E-06 |
| 10 min | 4.0E-11 | 3.2E-15 | 5.2E-26 | 1.6E-40 |
| 15 min | 1.5E-05 | 2.3E-139 | 3.8E-26 | 0.0E+00 |
| 20 min | 8.4E-01 | 9.7E-15 | 5.9E-09 | 4.3E-23 |
| 25 min | 2.0E-01 | 2.1E-05 | 2.0E-10 | 8.5E-25 |
| 30 min | 1.0E+00 | 4.8E-11 | 6.4E-12 | 2.2E-21 |
| 35 min | 7.1E-02 | 5.4E-05 | 2.8E-15 | 5.9E-23 |
| 40 min | 5.0E-01 | 6.0E-03 | 1.1E-11 | 1.8E-21 |
| 45 min | 9.0E-01 | 2.4E-05 | 5.1E-07 | 1.6E-22 |
| 50 min | 1.0E+00 | 6.2E-18 | 3.7E-03 | 6.2E-37 |
| 55 min | 7.1E-02 | 8.3E-32 | 1.6E-06 | 2.1E-64 |

**Table S3: p-value from ks-test comparing different time points.**

| | | | | | | |
|---|---|---|---|---|---|---|
| Gene | STL1 | | | | | |
| Conndition | 0.2M | | | | | |
| Compartment | Nucleus | | | | | |
| Replica | 1 | | | | | |
| timepoints | 0 min | 1 min | 2 min | 4 min | 6 min | 8 min |
| 0 min | 1.0E+00 | 1.0E+00 | 1.0E+00 | 1.0E+00 | 1.5E-11 | 4.8E-81 |
| 1 min | 1.0E+00 | 1.0E+00 | 1.0E+00 | 1.0E+00 | 1.5E-11 | 4.8E-81 |
| 2 min | 1.0E+00 | 1.0E+00 | 1.0E+00 | 1.0E+00 | 1.0E-09 | 4.1E-65 |
| 4 min | 1.0E+00 | 1.0E+00 | 1.0E+00 | 1.0E+00 | 5.1E-11 | 2.8E-76 |
| 6 min | 1.5E-11 | 1.5E-11 | 1.0E-09 | 5.1E-11 | 1.0E+00 | 2.2E-22 |
| 8 min | 4.8E-81 | 4.8E-81 | 4.1E-65 | 2.8E-76 | 2.2E-22 | 1.0E+00 |
| 10 min | 7.2E-74 | 7.2E-74 | 1.0E-63 | 5.6E-71 | 6.8E-27 | 3.9E-03 |
| 15 min | 4.0E-104 | 4.0E-104 | 3.0E-86 | 6.8E-99 | 6.2E-38 | 1.9E-06 |
| 20 min | 1.6E-31 | 1.6E-31 | 2.2E-25 | 1.1E-29 | 3.9E-03 | 5.4E-12 |
| 25 min | 1.7E-02 | 1.7E-02 | 3.4E-02 | 2.1E-02 | 8.5E-03 | 8.6E-37 |
| 30 min | 1.0E+00 | 1.0E+00 | 1.0E+00 | 1.0E+00 | 4.8E-09 | 2.2E-65 |
| 35 min | 1.0E+00 | 1.0E+00 | 1.0E+00 | 1.0E+00 | 1.4E-06 | 2.8E-41 |
| 40 min | 1.0E+00 | 1.0E+00 | 1.0E+00 | 1.0E+00 | 3.3E-09 | 1.0E-61 |
| 45 min | 1.0E+00 | 1.0E+00 | 1.0E+00 | 1.0E+00 | 4.7E-11 | 1.3E-76 |
| 50 min | 1.0E+00 | 1.0E+00 | 1.0E+00 | 1.0E+00 | 2.0E-07 | 8.1E-47 |
| 55 min | 1.0E+00 | 1.0E+00 | 1.0E+00 | 1.0E+00 | 2.6E-11 | 2.4E-80 |

| | | | | | | |
|---|---|---|---|---|---|---|
| Gene | STL1 | | | | | |
| Conndition | 0.2M | | | | | |
| Compartment | Nucleus | | | | | |
| Replica | 2 | | | | | |
| timepoints | 0 min | 1 min | 2 min | 4 min | 6 min | 8 min |
| 0 min | 1.0E+00 | 1.0E+00 | 1.0E+00 | 7.4E-01 | 2.7E-09 | 8.5E-28 |
| 1 min | 1.0E+00 | 1.0E+00 | 1.0E+00 | 7.4E-01 | 2.7E-09 | 8.5E-28 |
| 2 min | 1.0E+00 | 1.0E+00 | 1.0E+00 | 8.8E-01 | 8.8E-09 | 6.5E-27 |
| 4 min | 7.4E-01 | 7.4E-01 | 8.8E-01 | 1.0E+00 | 6.6E-06 | 1.2E-21 |
| 6 min | 2.7E-09 | 2.7E-09 | 8.8E-09 | 6.6E-06 | 1.0E+00 | 1.0E-05 |
| 8 min | 8.5E-28 | 8.5E-28 | 6.5E-27 | 1.2E-21 | 1.0E-05 | 1.0E+00 |
| 10 min | 3.5E-34 | 3.5E-34 | 3.5E-33 | 3.4E-27 | 1.9E-08 | 8.2E-01 |
| 15 min | 5.6E-51 | 5.6E-51 | 9.3E-50 | 2.4E-42 | 1.7E-17 | 1.5E-03 |
| 20 min | 2.8E-11 | 2.8E-11 | 1.0E-10 | 1.7E-07 | 1.0E+00 | 2.3E-04 |
| 25 min | 1.8E-01 | 1.8E-01 | 2.3E-01 | 8.0E-01 | 2.4E-01 | 1.1E-06 |
| 30 min | 1.0E+00 | 1.0E+00 | 1.0E+00 | 1.0E+00 | 1.3E-04 | 3.2E-14 |
| 35 min | 1.0E+00 | 1.0E+00 | 1.0E+00 | 8.4E-01 | 6.0E-09 | 3.3E-27 |
| 40 min | 1.0E+00 | 1.0E+00 | 1.0E+00 | 7.7E-01 | 8.9E-09 | 2.9E-26 |
| 45 min | 1.0E+00 | 1.0E+00 | 1.0E+00 | 8.6E-01 | 2.0E-07 | 3.3E-22 |
| 50 min | 1.0E+00 | 1.0E+00 | 1.0E+00 | 1.0E+00 | 6.8E-03 | 3.2E-08 |

| | | | | | | |
|---|---|---|---|---|---|---|
| 55 min | 1.0E+00 | 1.0E+00 | 1.0E+00 | 7.4E-01 | 2.7E-09 | 8.5E-28 |

| | | | | | | |
|---|---|---|---|---|---|---|
| Gene | STL1 | | | | | |
| Conndition | 0.2M | | | | | |
| Compartment | Cytoplasm | | | | | |
| Replica | 1 | | | | | |
| timepoints | 0 min | 1 min | 2 min | 4 min | 6 min | 8 min |
| 0 min | 1.0E+00 | 1.0E+00 | 1.0E+00 | 1.0E+00 | 3.7E-43 | 1.0E-264 |
| 1 min | 1.0E+00 | 1.0E+00 | 1.0E+00 | 1.0E+00 | 1.5E-44 | 1.4E-268 |
| 2 min | 1.0E+00 | 1.0E+00 | 1.0E+00 | 1.0E+00 | 6.6E-38 | 3.8E-217 |
| 4 min | 1.0E+00 | 1.0E+00 | 1.0E+00 | 1.0E+00 | 1.3E-42 | 3.5E-253 |
| 6 min | 3.7E-43 | 1.5E-44 | 6.6E-38 | 1.3E-42 | 1.0E+00 | 4.2E-78 |
| 8 min | 1.0E-264 | 1.4E-268 | 3.8E-217 | 3.5E-253 | 4.2E-78 | 1.0E+00 |
| 10 min | 6.7E-160 | 2.5E-162 | 5.2E-141 | 2.9E-156 | 4.0E-63 | 4.4E-08 |
| 15 min | 5.3E-281 | 1.1E-284 | 3.4E-237 | 9.2E-271 | 1.3E-112 | 8.0E-08 |
| 20 min | 5.1E-252 | 8.5E-256 | 7.8E-207 | 3.9E-241 | 1.8E-68 | 1.4E-20 |
| 25 min | 1.8E-53 | 6.4E-55 | 1.2E-47 | 7.9E-53 | 7.0E-02 | 9.7E-76 |
| 30 min | 5.4E-10 | 1.1E-10 | 3.3E-09 | 3.0E-10 | 2.5E-11 | 6.3E-148 |
| 35 min | 1.0E+00 | 9.9E-01 | 9.8E-01 | 9.9E-01 | 7.9E-22 | 1.5E-128 |
| 40 min | 1.0E+00 | 9.9E-01 | 9.8E-01 | 9.8E-01 | 2.0E-31 | 9.4E-194 |
| 45 min | 1.0E+00 | 9.9E-01 | 9.9E-01 | 9.9E-01 | 6.9E-38 | 9.4E-241 |
| 50 min | 1.0E+00 | 1.0E+00 | 1.0E+00 | 1.0E+00 | 4.4E-25 | 1.1E-146 |
| 55 min | 1.0E+00 | 1.0E+00 | 1.0E+00 | 1.0E+00 | 4.2E-41 | 5.7E-259 |

| | | | | | | |
|---|---|---|---|---|---|---|
| Gene | STL1 | | | | | |
| Conndition | 0.2M | | | | | |
| Compartment | Cytoplasm | | | | | |
| Replica | 2 | | | | | |
| timepoints | 0 min | 1 min | 2 min | 4 min | 6 min | 8 min |
| 0 min | 1.0E+00 | 1.0E+00 | 1.0E+00 | 1.1E-01 | 1.4E-24 | 1.8E-72 |
| 1 min | 1.0E+00 | 1.0E+00 | 1.0E+00 | 2.9E-02 | 6.8E-27 | 2.2E-76 |
| 2 min | 1.0E+00 | 1.0E+00 | 1.0E+00 | 5.7E-02 | 1.0E-25 | 2.1E-74 |
| 4 min | 1.1E-01 | 2.9E-02 | 5.7E-02 | 1.0E+00 | 9.2E-15 | 7.6E-55 |
| 6 min | 1.4E-24 | 6.8E-27 | 1.0E-25 | 9.2E-15 | 1.0E+00 | 6.1E-15 |
| 8 min | 1.8E-72 | 2.2E-76 | 2.1E-74 | 7.6E-55 | 6.1E-15 | 1.0E+00 |
| 10 min | 4.3E-108 | 6.0E-113 | 1.7E-110 | 5.7E-86 | 5.3E-45 | 1.8E-10 |
| 15 min | 7.7E-127 | 4.3E-132 | 1.9E-129 | 4.3E-108 | 8.9E-62 | 5.5E-19 |
| 20 min | 6.0E-113 | 6.6E-118 | 2.1E-115 | 2.7E-90 | 7.6E-33 | 2.9E-04 |
| 25 min | 1.5E-11 | 1.3E-12 | 4.4E-12 | 4.1E-07 | 6.1E-01 | 1.5E-09 |
| 30 min | 1.9E-02 | 6.0E-03 | 1.1E-02 | 8.0E-01 | 4.7E-05 | 1.6E-23 |
| 35 min | 6.4E-01 | 2.8E-01 | 4.4E-01 | 9.7E-01 | 3.1E-18 | 2.5E-61 |
| 40 min | 1.0E+00 | 1.0E+00 | 1.0E+00 | 1.5E-01 | 8.2E-23 | 8.3E-68 |
| 45 min | 1.0E+00 | 1.0E+00 | 1.0E+00 | 1.1E-01 | 1.3E-20 | 1.3E-59 |
| 50 min | 1.0E+00 | 1.0E+00 | 1.0E+00 | 6.8E-01 | 1.6E-07 | 1.3E-21 |
| 55 min | 1.0E+00 | 1.0E+00 | 1.0E+00 | 1.2E-01 | 2.8E-24 | 5.6E-72 |

| | | | | | | |
|---|---|---|---|---|---|---|
| Gene | STL1 | | | | | |
| Conndition | 0.4M | | | | | |
| Compartment | Cytoplasm | | | | | |
| Replica | 1 | | | | | |
| timepoints | 0 min | 1 min | 2 min | 4 min | 6 min | 8 min |
| 0 min | 1.0E+00 | 1.0E+00 | 1.0E+00 | 1.0E+00 | 1.6E-02 | 5.8E-28 |
| 1 min | 1.0E+00 | 1.0E+00 | 1.0E+00 | 1.0E+00 | 1.6E-02 | 5.8E-28 |
| 2 min | 1.0E+00 | 1.0E+00 | 1.0E+00 | 1.0E+00 | 2.4E-02 | 2.5E-27 |
| 4 min | 1.0E+00 | 1.0E+00 | 1.0E+00 | 1.0E+00 | 8.6E-02 | 3.8E-25 |
| 6 min | 1.6E-02 | 1.6E-02 | 2.4E-02 | 8.6E-02 | 1.0E+00 | 6.3E-15 |
| 8 min | 5.8E-28 | 5.8E-28 | 2.5E-27 | 3.8E-25 | 6.3E-15 | 1.0E+00 |
| 10 min | 1.8E-66 | 1.8E-66 | 1.8E-65 | 4.9E-62 | 4.8E-45 | 1.7E-08 |
| 15 min | 2.3E-102 | 2.3E-102 | 4.0E-101 | 7.5E-97 | 3.0E-75 | 2.3E-29 |
| 20 min | 3.8E-140 | 3.8E-140 | 1.1E-138 | 1.1E-133 | 4.9E-108 | 5.9E-53 |
| 25 min | 2.3E-102 | 2.3E-102 | 4.0E-101 | 7.5E-97 | 3.0E-75 | 1.8E-23 |
| 30 min | 1.9E-40 | 1.9E-40 | 1.0E-39 | 3.1E-37 | 4.1E-25 | 2.6E-02 |
| 35 min | 6.6E-03 | 6.6E-03 | 9.5E-03 | 3.1E-02 | 1.0E+00 | 3.9E-08 |
| 40 min | 9.1E-01 | 9.1E-01 | 9.7E-01 | 1.0E+00 | 2.8E-01 | 9.0E-23 |
| 45 min | 1.0E+00 | 1.0E+00 | 1.0E+00 | 1.0E+00 | 4.3E-02 | 2.2E-26 |
| 50 min | 1.0E+00 | 1.0E+00 | 1.0E+00 | 1.0E+00 | 2.3E-02 | 7.9E-27 |
| 55 min | 1.0E+00 | 1.0E+00 | 1.0E+00 | 1.0E+00 | 2.9E-02 | 5.2E-27 |

| | | | | | | |
|---|---|---|---|---|---|---|
| Gene | STL1 | | | | | |
| Conndition | 0.4M | | | | | |
| Compartment | Cytoplasm | | | | | |
| Replica | 2 | | | | | |
| timepoints | 0 min | 1 min | 2 min | 4 min | 6 min | 8 min |
| 0 min | 1.0E+00 | 1.0E+00 | 1.0E+00 | 1.0E+00 | 1.0E-01 | 2.3E-05 |
| 1 min | 1.0E+00 | 1.0E+00 | 1.0E+00 | 1.0E+00 | 7.0E-02 | 7.9E-06 |
| 2 min | 1.0E+00 | 1.0E+00 | 1.0E+00 | 1.0E+00 | 1.4E-01 | 5.8E-05 |
| 4 min | 1.0E+00 | 1.0E+00 | 1.0E+00 | 1.0E+00 | 3.1E-01 | 7.9E-04 |
| 6 min | 1.0E-01 | 7.0E-02 | 1.4E-01 | 3.1E-01 | 1.0E+00 | 4.6E-02 |
| 8 min | 2.3E-05 | 7.9E-06 | 5.8E-05 | 7.9E-04 | 4.6E-02 | 1.0E+00 |
| 10 min | 3.6E-23 | 7.2E-26 | 5.9E-21 | 4.7E-15 | 1.9E-15 | 5.5E-05 |
| 15 min | 1.0E-44 | 3.6E-49 | 5.7E-41 | 2.0E-30 | 2.7E-35 | 4.2E-17 |
| 20 min | 2.5E-78 | 3.5E-88 | 2.3E-70 | 1.5E-49 | 4.8E-67 | 4.9E-34 |
| 25 min | 2.0E-20 | 2.7E-21 | 1.3E-19 | 7.3E-17 | 2.7E-15 | 2.0E-08 |
| 30 min | 9.1E-67 | 8.6E-74 | 6.7E-61 | 1.2E-44 | 4.3E-56 | 1.2E-29 |
| 35 min | 3.1E-21 | 1.8E-23 | 2.2E-19 | 3.0E-14 | 4.9E-14 | 1.2E-04 |
| 40 min | 1.6E-04 | 5.6E-05 | 3.8E-04 | 4.1E-03 | 2.0E-01 | 1.0E+00 |
| 45 min | 1.0E+00 | 1.0E+00 | 1.0E+00 | 1.0E+00 | 4.0E-01 | 1.4E-03 |
| 50 min | 1.0E+00 | 1.0E+00 | 1.0E+00 | 1.0E+00 | 1.2E-01 | 1.8E-05 |
| 55 min | 1.0E+00 | 1.0E+00 | 1.0E+00 | 1.0E+00 | 1.2E-01 | 1.8E-05 |

| | | | | | | |
|---|---|---|---|---|---|---|
| Gene | STL1 | | | | | |
| Conndition | 0.4M | | | | | |
| Compartment | Cytoplasm | | | | | |
| Replica | 1 | | | | | |
| timepoints | 0 min | 1 min | 2 min | 4 min | 6 min | 8 min |
| 0 min | 1.0E+00 | 1.0E+00 | 1.0E+00 | 3.6E-01 | 1.0E-35 | 1.2E-108 |
| 1 min | 1.0E+00 | 1.0E+00 | 1.0E+00 | 4.5E-01 | 5.4E-35 | 2.2E-107 |
| 2 min | 1.0E+00 | 1.0E+00 | 1.0E+00 | 5.6E-01 | 2.8E-34 | 4.0E-106 |
| 4 min | 3.6E-01 | 4.5E-01 | 5.6E-01 | 1.0E+00 | 3.2E-26 | 1.5E-91 |
| 6 min | 1.0E-35 | 5.4E-35 | 2.8E-34 | 3.2E-26 | 1.0E+00 | 7.7E-35 |
| 8 min | 1.2E-108 | 2.2E-107 | 4.0E-106 | 1.5E-91 | 7.7E-35 | 1.0E+00 |
| 10 min | 3.4E-165 | 1.3E-163 | 4.7E-162 | 5.0E-146 | 4.0E-101 | 1.2E-24 |
| 15 min | 7.8E-163 | 2.9E-161 | 1.0E-159 | 1.6E-147 | 4.7E-128 | 2.1E-71 |
| 20 min | 5.1E-194 | 2.6E-192 | 1.9E-191 | 1.0E-184 | 2.6E-154 | 1.0E-74 |
| 25 min | 4.6E-188 | 2.2E-186 | 1.0E-184 | 2.2E-172 | 4.5E-143 | 8.0E-72 |
| 30 min | 1.9E-147 | 4.7E-146 | 1.2E-144 | 1.7E-128 | 2.1E-60 | 1.6E-07 |
| 35 min | 4.6E-62 | 2.9E-61 | 1.8E-60 | 2.3E-51 | 5.4E-10 | 4.8E-17 |
| 40 min | 9.8E-19 | 3.3E-18 | 1.1E-17 | 4.2E-12 | 6.2E-05 | 5.9E-57 |
| 45 min | 1.5E-04 | 2.6E-04 | 4.6E-04 | 8.6E-02 | 1.0E-15 | 1.3E-77 |
| 50 min | 5.4E-01 | 6.5E-01 | 7.6E-01 | 9.9E-01 | 1.7E-26 | 8.0E-91 |
| 55 min | 5.6E-01 | 6.7E-01 | 7.8E-01 | 1.0E+00 | 1.7E-27 | 6.7E-94 |

| | | | | | | |
|---|---|---|---|---|---|---|
| Gene | STL1 | | | | | |
| Conndition | 0.4M | | | | | |
| Compartment | Cytoplasm | | | | | |
| Replica | 2 | | | | | |
| timepoints | 0 min | 1 min | 2 min | 4 min | 6 min | 8 min |
| 0 min | 1.0E+00 | 1.0E+00 | 1.0E+00 | 1.0E+00 | 3.4E-19 | 5.3E-45 |
| 1 min | 1.0E+00 | 1.0E+00 | 1.0E+00 | 1.0E+00 | 8.4E-21 | 3.4E-48 |
| 2 min | 1.0E+00 | 1.0E+00 | 1.0E+00 | 1.0E+00 | 3.2E-17 | 2.5E-41 |
| 4 min | 1.0E+00 | 1.0E+00 | 1.0E+00 | 1.0E+00 | 7.5E-11 | 5.1E-29 |
| 6 min | 3.4E-19 | 8.4E-21 | 3.2E-17 | 7.5E-11 | 1.0E+00 | 2.0E-09 |
| 8 min | 5.3E-45 | 3.4E-48 | 2.5E-41 | 5.1E-29 | 2.0E-09 | 1.0E+00 |
| 10 min | 2.5E-91 | 1.8E-100 | 2.3E-82 | 8.8E-56 | 3.9E-48 | 1.0E-21 |
| 15 min | 6.1E-108 | 4.2E-117 | 1.1E-98 | 6.8E-70 | 2.4E-80 | 2.7E-51 |
| 20 min | 5.1E-163 | 2.7E-181 | 3.1E-146 | 1.1E-98 | 2.5E-144 | 2.6E-96 |
| 25 min | 1.2E-58 | 2.1E-60 | 3.0E-56 | 1.7E-47 | 2.7E-45 | 6.5E-35 |
| 30 min | 2.6E-125 | 1.0E-136 | 4.0E-114 | 6.1E-81 | 6.2E-102 | 4.9E-64 |
| 35 min | 3.0E-132 | 1.7E-144 | 2.5E-120 | 2.9E-84 | 5.3E-79 | 4.2E-32 |
| 40 min | 1.8E-81 | 5.3E-89 | 7.2E-74 | 1.1E-50 | 3.9E-27 | 2.8E-03 |
| 45 min | 8.6E-01 | 9.1E-01 | 8.9E-01 | 1.0E+00 | 2.2E-08 | 5.3E-25 |
| 50 min | 1.2E-03 | 9.4E-04 | 2.6E-03 | 6.6E-02 | 1.2E-07 | 7.7E-29 |
| 55 min | 6.9E-01 | 7.7E-01 | 7.7E-01 | 1.0E+00 | 1.3E-15 | 4.5E-41 |

| | | | | | | |
|---|---|---|---|---|---|---|
| Gene | CTT1 | | | | | |
| Conndition | 0.2M | | | | | |
| Compartment | Nucleus | | | | | |
| Replica | 1 | | | | | |
| timepoints | 0 min | 1 min | 2 min | 4 min | 6 min | 8 min |
| 0 min | 1.0E+00 | 1.0E+00 | 1.0E+00 | 8.2E-01 | 4.0E-18 | 4.8E-81 |
| 1 min | 1.0E+00 | 1.0E+00 | 1.0E+00 | 1.0E+00 | 7.9E-16 | 1.8E-75 |
| 2 min | 1.0E+00 | 1.0E+00 | 1.0E+00 | 9.2E-01 | 4.2E-15 | 7.8E-65 |
| 4 min | 8.2E-01 | 1.0E+00 | 9.2E-01 | 1.0E+00 | 3.2E-13 | 2.3E-66 |
| 6 min | 4.0E-18 | 7.9E-16 | 4.2E-15 | 3.2E-13 | 1.0E+00 | 5.8E-15 |
| 8 min | 4.8E-81 | 1.8E-75 | 7.8E-65 | 2.3E-66 | 5.8E-15 | 1.0E+00 |
| 10 min | 1.1E-55 | 5.8E-52 | 7.3E-48 | 9.5E-47 | 1.0E-11 | 9.9E-01 |
| 15 min | 7.8E-77 | 7.9E-72 | 2.1E-63 | 5.2E-64 | 2.0E-16 | 2.9E-02 |
| 20 min | 6.8E-49 | 1.4E-44 | 4.3E-39 | 2.0E-38 | 1.8E-04 | 1.7E-04 |
| 25 min | 1.7E-43 | 4.0E-40 | 4.5E-37 | 1.3E-35 | 2.2E-06 | 1.8E-01 |
| 30 min | 4.7E-30 | 5.7E-27 | 8.7E-25 | 4.4E-23 | 1.3E-01 | 2.4E-08 |
| 35 min | 1.3E-06 | 1.6E-05 | 7.4E-06 | 1.7E-04 | 4.0E-01 | 5.4E-16 |
| 40 min | 3.0E-09 | 1.1E-07 | 9.0E-08 | 4.2E-06 | 1.6E-01 | 2.7E-24 |
| 45 min | 3.1E-15 | 5.8E-13 | 2.5E-12 | 1.9E-10 | 7.1E-01 | 3.2E-24 |
| 50 min | 9.2E-08 | 1.6E-06 | 8.7E-07 | 2.6E-05 | 4.4E-01 | 3.0E-17 |
| 55 min | 9.4E-32 | 2.6E-28 | 2.2E-25 | 7.4E-24 | 4.7E-01 | 7.3E-12 |

| | | | | | | |
|---|---|---|---|---|---|---|
| Gene | CTT1 | | | | | |
| Conndition | 0.2M | | | | | |
| Compartment | Nucleus | | | | | |
| Replica | 2 | | | | | |
| timepoints | 0 min | 1 min | 2 min | 4 min | 6 min | 8 min |
| 0 min | 1.0E+00 | 1.0E+00 | 1.0E+00 | 9.0E-03 | 5.3E-24 | 3.5E-68 |
| 1 min | 1.0E+00 | 1.0E+00 | 1.0E+00 | 9.0E-03 | 5.3E-24 | 3.5E-68 |
| 2 min | 1.0E+00 | 1.0E+00 | 1.0E+00 | 9.0E-03 | 5.3E-24 | 3.5E-68 |
| 4 min | 9.0E-03 | 9.0E-03 | 9.0E-03 | 1.0E+00 | 1.8E-11 | 1.3E-45 |
| 6 min | 5.3E-24 | 5.3E-24 | 5.3E-24 | 1.8E-11 | 1.0E+00 | 2.6E-12 |
| 8 min | 3.5E-68 | 3.5E-68 | 3.5E-68 | 1.3E-45 | 2.6E-12 | 1.0E+00 |
| 10 min | 5.3E-24 | 5.3E-24 | 5.3E-24 | 1.8E-11 | 1.0E+00 | 2.6E-12 |
| 15 min | 4.7E-53 | 4.7E-53 | 4.7E-53 | 3.5E-33 | 3.5E-06 | 1.3E-01 |
| 20 min | 3.5E-34 | 3.5E-34 | 3.5E-34 | 1.0E-18 | 2.5E-01 | 1.0E-06 |
| 25 min | 1.3E-12 | 1.3E-12 | 1.3E-12 | 1.6E-06 | 7.1E-01 | 1.1E-04 |
| 30 min | 3.3E-04 | 3.3E-04 | 3.3E-04 | 4.1E-01 | 6.1E-03 | 3.1E-17 |
| 35 min | 7.1E-05 | 7.1E-05 | 7.1E-05 | 8.4E-01 | 5.8E-08 | 2.8E-38 |
| 40 min | 3.0E-03 | 3.0E-03 | 3.0E-03 | 1.0E+00 | 1.3E-09 | 1.3E-40 |
| 45 min | 1.9E-03 | 1.9E-03 | 1.9E-03 | 1.0E+00 | 4.4E-07 | 4.5E-32 |
| 50 min | 1.9E-01 | 1.9E-01 | 1.9E-01 | 1.0E+00 | 7.7E-03 | 6.5E-12 |
| 55 min | 8.3E-08 | 8.3E-08 | 8.3E-08 | 7.9E-02 | 5.3E-05 | 2.7E-31 |

| | | | | | | |
|---|---|---|---|---|---|---|
| Gene | CTT1 | | | | | |
| Conndition | 0.2M | | | | | |
| Compartment | Cytoplasm | | | | | |
| Replica | 1 | | | | | |
| timepoints | 0 min | 1 min | 2 min | 4 min | 6 min | 8 min |
| 0 min | 1.0E+00 | 5.3E-01 | 1.0E+00 | 1.7E-08 | 1.0E-21 | 3.6E-245 |
| 1 min | 5.3E-01 | 1.0E+00 | 9.9E-01 | 6.8E-05 | 3.8E-20 | 4.4E-242 |
| 2 min | 1.0E+00 | 9.9E-01 | 1.0E+00 | 1.1E-05 | 1.0E-17 | 4.6E-197 |
| 4 min | 1.7E-08 | 6.8E-05 | 1.1E-05 | 1.0E+00 | 2.0E-12 | 1.6E-199 |
| 6 min | 1.0E-21 | 3.8E-20 | 1.0E-17 | 2.0E-12 | 1.0E+00 | 1.3E-89 |
| 8 min | 3.6E-245 | 4.4E-242 | 4.6E-197 | 1.6E-199 | 1.3E-89 | 1.0E+00 |
| 10 min | 2.1E-153 | 8.9E-151 | 6.8E-132 | 6.4E-128 | 1.3E-61 | 5.4E-02 |
| 15 min | 1.9E-233 | 8.1E-234 | 3.7E-196 | 3.5E-196 | 9.2E-95 | 1.1E-01 |
| 20 min | 2.1E-290 | 2.1E-290 | 4.3E-236 | 3.5E-242 | 2.2E-116 | 2.6E-03 |
| 25 min | 9.6E-195 | 1.1E-192 | 5.7E-167 | 2.6E-163 | 4.8E-81 | 9.7E-04 |
| 30 min | 2.5E-249 | 2.3E-240 | 1.2E-204 | 4.7E-202 | 1.5E-100 | 1.3E-08 |
| 35 min | 2.8E-98 | 2.7E-93 | 2.0E-86 | 7.5E-71 | 2.3E-34 | 2.2E-28 |
| 40 min | 4.1E-110 | 1.1E-103 | 1.4E-91 | 2.2E-74 | 1.0E-38 | 2.9E-38 |
| 45 min | 1.5E-137 | 1.4E-129 | 2.7E-110 | 4.1E-92 | 5.5E-42 | 3.4E-47 |
| 50 min | 5.4E-58 | 2.7E-53 | 1.6E-49 | 4.7E-37 | 1.2E-17 | 8.9E-33 |
| 55 min | 3.8E-162 | 3.2E-153 | 1.2E-128 | 2.4E-110 | 1.1E-57 | 8.9E-34 |

| | | | | | | |
|---|---|---|---|---|---|---|
| Gene | CTT1 | | | | | |
| Conndition | 0.2M | | | | | |
| Compartment | Cytoplasm | | | | | |
| Replica | 2 | | | | | |
| timepoints | 0 min | 1 min | 2 min | 4 min | 6 min | 8 min |
| 0 min | 1.0E+00 | 8.8E-01 | 1.0E+00 | 9.2E-02 | 1.8E-43 | 3.0E-108 |
| 1 min | 8.8E-01 | 1.0E+00 | 1.0E+00 | 2.5E-03 | 5.6E-51 | 1.1E-116 |
| 2 min | 1.0E+00 | 1.0E+00 | 1.0E+00 | 3.4E-02 | 8.9E-46 | 1.9E-115 |
| 4 min | 9.2E-02 | 2.5E-03 | 3.4E-02 | 1.0E+00 | 7.2E-32 | 6.0E-96 |
| 6 min | 1.8E-43 | 5.6E-51 | 8.9E-46 | 7.2E-32 | 1.0E+00 | 1.7E-24 |
| 8 min | 3.0E-108 | 1.1E-116 | 1.9E-115 | 6.0E-96 | 1.7E-24 | 1.0E+00 |
| 10 min | 1.2E-116 | 3.0E-124 | 5.7E-123 | 6.2E-104 | 8.6E-28 | 5.6E-01 |
| 15 min | 1.5E-141 | 3.4E-149 | 2.2E-146 | 1.7E-127 | 1.0E-42 | 7.1E-07 |
| 20 min | 6.7E-163 | 4.2E-171 | 4.3E-168 | 8.6E-148 | 6.7E-54 | 6.2E-10 |
| 25 min | 1.0E-46 | 1.8E-50 | 6.3E-50 | 3.4E-41 | 5.5E-10 | 5.5E-02 |
| 30 min | 2.2E-56 | 7.3E-61 | 3.3E-60 | 2.2E-49 | 8.1E-10 | 5.9E-04 |
| 35 min | 5.7E-86 | 1.3E-93 | 1.7E-92 | 4.1E-74 | 2.7E-09 | 1.7E-21 |
| 40 min | 1.2E-55 | 7.7E-64 | 3.6E-58 | 3.2E-40 | 6.7E-04 | 4.3E-38 |
| 45 min | 1.5E-41 | 4.8E-48 | 1.5E-43 | 2.0E-29 | 1.4E-02 | 6.9E-30 |
| 50 min | 3.9E-11 | 4.1E-13 | 9.7E-12 | 3.8E-08 | 1.3E-01 | 4.6E-13 |
| 55 min | 2.5E-57 | 6.4E-66 | 3.1E-62 | 2.4E-47 | 5.7E-02 | 4.5E-22 |

| | |
|---|---|
| Gene | CTT1 |

| | | | | | | |
|---|---|---|---|---|---|---|
| Conndition | 0.4M | | | | | |
| Compartment | Nucleus | | | | | |
| Replica | 1 | | | | | |
| timepoints | 0 min | 1 min | 2 min | 4 min | 6 min | 8 min |
| 0 min | 1.0E+00 | 1.0E+00 | 1.0E+00 | 9.7E-01 | 6.2E-08 | 1.3E-34 |
| 1 min | 1.0E+00 | 1.0E+00 | 1.0E+00 | 6.7E-01 | 3.7E-09 | 3.7E-37 |
| 2 min | 1.0E+00 | 1.0E+00 | 1.0E+00 | 9.4E-01 | 4.2E-08 | 5.8E-35 |
| 4 min | 9.7E-01 | 6.7E-01 | 9.4E-01 | 1.0E+00 | 1.3E-05 | 1.9E-29 |
| 6 min | 6.2E-08 | 3.7E-09 | 4.2E-08 | 1.3E-05 | 1.0E+00 | 3.7E-10 |
| 8 min | 1.3E-34 | 3.7E-37 | 5.8E-35 | 1.9E-29 | 3.7E-10 | 1.0E+00 |
| 10 min | 5.6E-68 | 1.5E-71 | 1.8E-68 | 1.3E-60 | 1.9E-30 | 2.2E-06 |
| 15 min | 1.2E-92 | 7.5E-97 | 2.9E-93 | 5.2E-84 | 1.5E-47 | 1.6E-14 |
| 20 min | 1.6E-150 | 7.5E-156 | 2.8E-151 | 2.0E-139 | 6.7E-91 | 4.8E-49 |
| 25 min | 1.9E-94 | 1.1E-98 | 4.8E-95 | 1.0E-85 | 8.0E-49 | 6.8E-15 |
| 30 min | 7.5E-145 | 9.4E-150 | 1.5E-145 | 1.3E-134 | 1.2E-89 | 3.0E-61 |
| 35 min | 3.6E-47 | 1.2E-49 | 1.6E-47 | 4.3E-42 | 2.6E-21 | 2.6E-04 |
| 40 min | 4.0E-47 | 4.1E-50 | 1.5E-47 | 5.0E-41 | 3.5E-17 | 9.8E-03 |
| 45 min | 4.8E-45 | 5.7E-48 | 1.9E-45 | 4.3E-39 | 6.0E-16 | 1.2E-02 |
| 50 min | 4.8E-22 | 4.6E-24 | 2.5E-22 | 5.0E-18 | 1.9E-04 | 1.0E-01 |
| 55 min | 1.3E-20 | 1.3E-22 | 6.7E-21 | 1.1E-16 | 1.7E-03 | 2.6E-02 |

| | | | | | | |
|---|---|---|---|---|---|---|
| Gene | CTT1 | | | | | |
| Conndition | 0.4M | | | | | |
| Compartment | Nucleus | | | | | |
| Replica | 2 | | | | | |
| timepoints | 0 min | 1 min | 2 min | 4 min | 6 min | 8 min |
| 0 min | 1.0E+00 | 1.0E+00 | 1.0E+00 | 1.0E+00 | 5.1E-06 | 1.3E-21 |
| 1 min | 1.0E+00 | 1.0E+00 | 1.0E+00 | 1.0E+00 | 5.4E-08 | 1.1E-25 |
| 2 min | 1.0E+00 | 1.0E+00 | 1.0E+00 | 1.0E+00 | 2.2E-06 | 1.6E-21 |
| 4 min | 1.0E+00 | 1.0E+00 | 1.0E+00 | 1.0E+00 | 2.2E-03 | 6.1E-14 |
| 6 min | 5.1E-06 | 5.4E-08 | 2.2E-06 | 2.2E-03 | 1.0E+00 | 2.2E-07 |
| 8 min | 1.3E-21 | 1.1E-25 | 1.6E-21 | 6.1E-14 | 2.2E-07 | 1.0E+00 |
| 10 min | 4.0E-46 | 6.0E-55 | 3.1E-44 | 3.5E-28 | 1.1E-22 | 8.4E-03 |
| 15 min | 5.5E-74 | 2.7E-85 | 8.7E-71 | 3.5E-48 | 4.6E-46 | 2.4E-13 |
| 20 min | 1.7E-97 | 7.4E-115 | 2.3E-91 | 3.5E-59 | 1.5E-63 | 9.8E-19 |
| 25 min | 3.2E-42 | 6.4E-46 | 3.7E-42 | 1.7E-33 | 1.2E-26 | 3.6E-13 |
| 30 min | 8.8E-73 | 1.9E-84 | 1.9E-69 | 1.3E-46 | 1.4E-44 | 9.7E-14 |
| 35 min | 4.0E-88 | 6.1E-102 | 1.1E-83 | 2.1E-56 | 2.8E-57 | 8.7E-27 |
| 40 min | 2.4E-75 | 7.0E-88 | 1.3E-71 | 1.8E-47 | 3.4E-46 | 6.3E-21 |
| 45 min | 1.2E-22 | 6.1E-26 | 9.7E-23 | 5.1E-16 | 1.6E-09 | 6.3E-01 |
| 50 min | 7.6E-30 | 2.1E-36 | 6.0E-29 | 1.2E-17 | 5.8E-11 | 3.8E-01 |
| 55 min | 1.7E-11 | 1.2E-14 | 1.1E-11 | 1.3E-06 | 1.8E-01 | 1.7E-03 |

| | |
|---|---|
| Gene | CTT1 |
| Conndition | 0.4M |

| | | | | | | |
|---|---|---|---|---|---|---|
| Compartment | Cytoplasm | | | | | |
| Replica | 1 | | | | | |
| timepoints | 0 min | 1 min | 2 min | 4 min | 6 min | 8 min |
| 0 min | 1.0E+00 | 1.1E-02 | 3.6E-01 | 1.4E-01 | 8.8E-24 | 2.1E-97 |
| 1 min | 1.1E-02 | 1.0E+00 | 7.2E-01 | 9.8E-01 | 3.3E-33 | 7.2E-104 |
| 2 min | 3.6E-01 | 7.2E-01 | 1.0E+00 | 1.0E+00 | 1.9E-28 | 3.0E-103 |
| 4 min | 1.4E-01 | 9.8E-01 | 1.0E+00 | 1.0E+00 | 1.9E-28 | 9.0E-101 |
| 6 min | 8.8E-24 | 3.3E-33 | 1.9E-28 | 1.9E-28 | 1.0E+00 | 1.9E-31 |
| 8 min | 2.1E-97 | 7.2E-104 | 3.0E-103 | 9.0E-101 | 1.9E-31 | 1.0E+00 |
| 10 min | 1.6E-150 | 1.6E-150 | 1.6E-150 | 1.5E-144 | 7.5E-94 | 8.0E-32 |
| 15 min | 9.3E-190 | 1.9E-191 | 6.5E-189 | 3.2E-187 | 8.6E-153 | 3.3E-87 |
| 20 min | 3.4E-199 | 4.7E-200 | 1.9E-197 | 1.9E-197 | 3.6E-169 | 2.9E-101 |
| 25 min | 1.3E-190 | 1.9E-191 | 6.5E-189 | 2.2E-186 | 3.4E-173 | 4.9E-125 |
| 30 min | 3.9E-168 | 6.9E-169 | 8.4E-168 | 2.0E-164 | 1.3E-129 | 1.2E-59 |
| 35 min | 5.1E-128 | 1.4E-128 | 7.2E-127 | 3.6E-124 | 1.2E-101 | 5.0E-59 |
| 40 min | 1.3E-163 | 1.3E-163 | 2.1E-164 | 2.2E-157 | 4.9E-108 | 5.8E-43 |
| 45 min | 4.2E-135 | 4.2E-135 | 4.2E-135 | 1.9E-129 | 1.6E-62 | 9.8E-14 |
| 50 min | 4.6E-96 | 4.2E-103 | 1.7E-102 | 4.7E-100 | 4.2E-31 | 3.2E-02 |
| 55 min | 1.9E-83 | 1.2E-92 | 4.5E-92 | 9.9E-90 | 2.9E-19 | 7.3E-04 |

| | | | | | | |
|---|---|---|---|---|---|---|
| Gene | CTT1 | | | | | |
| Conndition | 0.4M | | | | | |
| Compartment | Cytoplasm | | | | | |
| Replica | 2 | | | | | |
| timepoints | 0 min | 1 min | 2 min | 4 min | 6 min | 8 min |
| 0 min | 1.0E+00 | 5.0E-01 | 4.7E-01 | 1.5E-01 | 2.2E-10 | 7.9E-47 |
| 1 min | 5.0E-01 | 1.0E+00 | 7.3E-03 | 2.3E-03 | 2.5E-07 | 8.4E-45 |
| 2 min | 4.7E-01 | 7.3E-03 | 1.0E+00 | 6.5E-01 | 5.1E-13 | 6.6E-45 |
| 4 min | 1.5E-01 | 2.3E-03 | 6.5E-01 | 1.0E+00 | 7.5E-12 | 5.6E-40 |
| 6 min | 2.2E-10 | 2.5E-07 | 5.1E-13 | 7.5E-12 | 1.0E+00 | 2.4E-20 |
| 8 min | 7.9E-47 | 8.4E-45 | 6.6E-45 | 5.6E-40 | 2.4E-20 | 1.0E+00 |
| 10 min | 2.1E-92 | 2.2E-103 | 1.3E-86 | 2.2E-68 | 1.3E-60 | 5.1E-16 |
| 15 min | 8.9E-123 | 2.9E-135 | 5.7E-112 | 4.4E-87 | 7.5E-100 | 1.1E-55 |
| 20 min | 3.7E-174 | 7.6E-195 | 5.7E-154 | 3.6E-111 | 3.7E-177 | 2.5E-109 |
| 25 min | 2.5E-58 | 2.1E-60 | 4.6E-56 | 1.1E-50 | 1.6E-53 | 3.6E-39 |
| 30 min | 7.1E-141 | 9.0E-157 | 1.2E-127 | 4.4E-97 | 3.3E-140 | 3.6E-88 |
| 35 min | 5.3E-144 | 1.3E-159 | 1.1E-130 | 1.2E-99 | 2.4E-132 | 9.0E-80 |
| 40 min | 2.8E-136 | 1.6E-152 | 1.5E-122 | 3.2E-93 | 1.5E-114 | 4.6E-68 |
| 45 min | 1.1E-82 | 2.3E-88 | 4.5E-79 | 1.6E-67 | 2.3E-57 | 2.9E-17 |
| 50 min | 2.5E-119 | 4.0E-134 | 3.3E-111 | 7.9E-88 | 6.3E-82 | 4.0E-30 |
| 55 min | 1.4E-80 | 3.6E-84 | 1.4E-71 | 3.0E-61 | 3.4E-44 | 1.1E-06 |

| 10 min | 15 min | 20 min | 25 min | 30 min | 35 min | 40 min |
|---|---|---|---|---|---|---|
| 7.2E-74 | 4.0E-104 | 1.6E-31 | 1.7E-02 | 1.0E+00 | 1.0E+00 | 1.0E+00 |
| 7.2E-74 | 4.0E-104 | 1.6E-31 | 1.7E-02 | 1.0E+00 | 1.0E+00 | 1.0E+00 |
| 1.0E-63 | 3.0E-86 | 2.2E-25 | 3.4E-02 | 1.0E+00 | 1.0E+00 | 1.0E+00 |
| 5.6E-71 | 6.8E-99 | 1.1E-29 | 2.1E-02 | 1.0E+00 | 1.0E+00 | 1.0E+00 |
| 6.8E-27 | 6.2E-38 | 3.9E-03 | 8.5E-03 | 4.8E-09 | 1.4E-06 | 3.3E-09 |
| 3.9E-03 | 1.9E-06 | 5.4E-12 | 8.6E-37 | 2.2E-65 | 2.8E-41 | 1.0E-61 |
| 1.0E+00 | 7.5E-01 | 6.3E-17 | 2.6E-40 | 2.0E-63 | 2.4E-45 | 2.7E-61 |
| 7.5E-01 | 1.0E+00 | 6.0E-26 | 4.6E-54 | 8.2E-87 | 2.2E-57 | 2.5E-82 |
| 6.3E-17 | 6.0E-26 | 1.0E+00 | 1.1E-10 | 9.8E-25 | 4.6E-16 | 5.8E-24 |
| 2.6E-40 | 4.6E-54 | 1.1E-10 | 1.0E+00 | 7.9E-02 | 1.4E-01 | 4.6E-02 |
| 2.0E-63 | 8.2E-87 | 9.8E-25 | 7.9E-02 | 1.0E+00 | 1.0E+00 | 1.0E+00 |
| 2.4E-45 | 2.2E-57 | 4.6E-16 | 1.4E-01 | 1.0E+00 | 1.0E+00 | 1.0E+00 |
| 2.7E-61 | 2.5E-82 | 5.8E-24 | 4.6E-02 | 1.0E+00 | 1.0E+00 | 1.0E+00 |
| 3.5E-71 | 3.0E-99 | 8.2E-30 | 2.1E-02 | 1.0E+00 | 1.0E+00 | 1.0E+00 |
| 4.8E-50 | 2.7E-64 | 2.5E-18 | 8.6E-02 | 1.0E+00 | 1.0E+00 | 1.0E+00 |
| 2.5E-73 | 2.2E-103 | 4.3E-31 | 2.1E-02 | 1.0E+00 | 1.0E+00 | 1.0E+00 |

| 10 min | 15 min | 20 min | 25 min | 30 min | 35 min | 40 min |
|---|---|---|---|---|---|---|
| 3.5E-34 | 5.6E-51 | 2.8E-11 | 1.8E-01 | 1.0E+00 | 1.0E+00 | 1.0E+00 |
| 3.5E-34 | 5.6E-51 | 2.8E-11 | 1.8E-01 | 1.0E+00 | 1.0E+00 | 1.0E+00 |
| 3.5E-33 | 9.3E-50 | 1.0E-10 | 2.3E-01 | 1.0E+00 | 1.0E+00 | 1.0E+00 |
| 3.4E-27 | 2.4E-42 | 1.7E-07 | 8.0E-01 | 1.0E+00 | 8.4E-01 | 7.7E-01 |
| 1.9E-08 | 1.7E-17 | 1.0E+00 | 2.4E-01 | 1.3E-04 | 6.0E-09 | 8.9E-09 |
| 8.2E-01 | 1.5E-03 | 2.3E-04 | 1.1E-06 | 3.2E-14 | 3.3E-27 | 2.9E-26 |
| 1.0E+00 | 4.1E-02 | 9.5E-07 | 1.6E-08 | 2.8E-17 | 1.7E-33 | 3.1E-32 |
| 4.1E-02 | 1.0E+00 | 5.6E-15 | 3.2E-14 | 6.2E-26 | 3.7E-50 | 4.6E-48 |
| 9.5E-07 | 5.6E-15 | 1.0E+00 | 8.5E-02 | 1.3E-05 | 6.6E-11 | 1.2E-10 |
| 1.6E-08 | 3.2E-14 | 8.5E-02 | 1.0E+00 | 4.9E-01 | 2.1E-01 | 1.9E-01 |
| 2.8E-17 | 6.2E-26 | 1.3E-05 | 4.9E-01 | 1.0E+00 | 1.0E+00 | 1.0E+00 |
| 1.7E-33 | 3.7E-50 | 6.6E-11 | 2.1E-01 | 1.0E+00 | 1.0E+00 | 1.0E+00 |
| 3.1E-32 | 4.6E-48 | 1.2E-10 | 1.9E-01 | 1.0E+00 | 1.0E+00 | 1.0E+00 |
| 4.1E-27 | 2.1E-40 | 5.2E-09 | 2.3E-01 | 1.0E+00 | 1.0E+00 | 1.0E+00 |
| 7.8E-10 | 1.6E-14 | 1.9E-03 | 6.2E-01 | 1.0E+00 | 1.0E+00 | 1.0E+00 |

| | | | | | | |
|---|---|---|---|---|---|---|
| 3.5E-34 | 5.6E-51 | 2.8E-11 | 1.8E-01 | 1.0E+00 | 1.0E+00 | 1.0E+00 |

| 10 min | 15 min | 20 min | 25 min | 30 min | 35 min | 40 min |
|---|---|---|---|---|---|---|
| 6.7E-160 | 5.3E-281 | 5.1E-252 | 1.8E-53 | 5.4E-10 | 1.0E+00 | 1.0E+00 |
| 2.5E-162 | 1.1E-284 | 8.5E-256 | 6.4E-55 | 1.1E-10 | 9.9E-01 | 9.9E-01 |
| 5.2E-141 | 3.4E-237 | 7.8E-207 | 1.2E-47 | 3.3E-09 | 9.8E-01 | 9.8E-01 |
| 2.9E-156 | 9.2E-271 | 3.9E-241 | 7.9E-53 | 3.0E-10 | 9.9E-01 | 9.8E-01 |
| 4.0E-63 | 1.3E-112 | 1.8E-68 | 7.0E-02 | 2.5E-11 | 7.9E-22 | 2.0E-31 |
| 4.4E-08 | 8.0E-08 | 1.4E-20 | 9.7E-76 | 6.3E-148 | 1.5E-128 | 9.4E-194 |
| 1.0E+00 | 2.9E-06 | 3.1E-33 | 1.1E-71 | 1.9E-111 | 2.4E-93 | 3.0E-127 |
| 2.9E-06 | 1.0E+00 | 1.1E-30 | 1.6E-109 | 7.2E-186 | 2.0E-148 | 1.9E-214 |
| 3.1E-33 | 1.1E-30 | 1.0E+00 | 5.7E-53 | 4.0E-134 | 3.2E-122 | 3.0E-184 |
| 1.1E-71 | 1.6E-109 | 5.7E-53 | 1.0E+00 | 2.6E-18 | 3.0E-29 | 1.2E-40 |
| 1.9E-111 | 7.2E-186 | 4.0E-134 | 2.6E-18 | 1.0E+00 | 2.2E-04 | 1.8E-06 |
| 2.4E-93 | 2.0E-148 | 3.2E-122 | 3.0E-29 | 2.2E-04 | 1.0E+00 | 1.0E+00 |
| 3.0E-127 | 1.9E-214 | 3.0E-184 | 1.2E-40 | 1.8E-06 | 1.0E+00 | 1.0E+00 |
| 4.7E-148 | 8.0E-259 | 5.6E-229 | 6.8E-48 | 6.0E-08 | 1.0E+00 | 1.0E+00 |
| 7.0E-104 | 2.6E-167 | 1.6E-139 | 4.1E-33 | 2.9E-05 | 1.0E+00 | 1.0E+00 |
| 2.8E-156 | 1.5E-275 | 2.1E-246 | 2.4E-51 | 5.3E-09 | 1.0E+00 | 1.0E+00 |

| 10 min | 15 min | 20 min | 25 min | 30 min | 35 min | 40 min |
|---|---|---|---|---|---|---|
| 4.3E-108 | 7.7E-127 | 6.0E-113 | 1.5E-11 | 1.9E-02 | 6.4E-01 | 1.0E+00 |
| 6.0E-113 | 4.3E-132 | 6.6E-118 | 1.3E-12 | 6.0E-03 | 2.8E-01 | 1.0E+00 |
| 1.7E-110 | 1.9E-129 | 2.1E-115 | 4.4E-12 | 1.1E-02 | 4.4E-01 | 1.0E+00 |
| 5.7E-86 | 4.3E-108 | 2.7E-90 | 4.1E-07 | 8.0E-01 | 9.7E-01 | 1.5E-01 |
| 5.3E-45 | 8.9E-62 | 7.6E-33 | 6.1E-01 | 4.7E-05 | 3.1E-18 | 8.2E-23 |
| 1.8E-10 | 5.5E-19 | 2.9E-04 | 1.5E-09 | 1.6E-23 | 2.5E-61 | 8.3E-68 |
| 1.0E+00 | 9.2E-02 | 1.7E-17 | 4.8E-27 | 2.1E-42 | 3.5E-94 | 4.0E-101 |
| 9.2E-02 | 1.0E+00 | 4.4E-22 | 1.4E-35 | 5.2E-54 | 6.6E-118 | 1.0E-118 |
| 1.7E-17 | 4.4E-22 | 1.0E+00 | 1.4E-16 | 5.5E-40 | 1.0E-98 | 1.1E-105 |
| 4.8E-27 | 1.4E-35 | 1.4E-16 | 1.0E+00 | 2.1E-03 | 1.1E-08 | 4.2E-11 |
| 2.1E-42 | 5.2E-54 | 5.5E-40 | 2.1E-03 | 1.0E+00 | 2.9E-01 | 2.6E-02 |
| 3.5E-94 | 6.6E-118 | 1.0E-98 | 1.1E-08 | 2.9E-01 | 1.0E+00 | 7.4E-01 |
| 4.0E-101 | 1.0E-118 | 1.1E-105 | 4.2E-11 | 2.6E-02 | 7.4E-01 | 1.0E+00 |
| 7.2E-88 | 6.9E-103 | 9.3E-92 | 5.7E-11 | 1.8E-02 | 5.5E-01 | 1.0E+00 |
| 3.1E-31 | 1.6E-36 | 1.3E-32 | 6.3E-06 | 2.1E-01 | 9.8E-01 | 1.0E+00 |
| 1.7E-107 | 3.4E-126 | 2.5E-112 | 2.0E-11 | 2.2E-02 | 6.9E-01 | 1.0E+00 |

| 10 min | 15 min | 20 min | 25 min | 30 min | 35 min | 40 min |
|---|---|---|---|---|---|---|
| 1.8E-66 | 2.3E-102 | 3.8E-140 | 2.3E-102 | 1.9E-40 | 6.6E-03 | 9.1E-01 |
| 1.8E-66 | 2.3E-102 | 3.8E-140 | 2.3E-102 | 1.9E-40 | 6.6E-03 | 9.1E-01 |
| 1.8E-65 | 4.0E-101 | 1.1E-138 | 4.0E-101 | 1.0E-39 | 9.5E-03 | 9.7E-01 |
| 4.9E-62 | 7.5E-97 | 1.1E-133 | 7.5E-97 | 3.1E-37 | 3.1E-02 | 1.0E+00 |
| 4.8E-45 | 3.0E-75 | 4.9E-108 | 3.0E-75 | 4.1E-25 | 1.0E+00 | 2.8E-01 |
| 1.7E-08 | 2.3E-29 | 5.9E-53 | 1.8E-23 | 2.6E-02 | 3.9E-08 | 9.0E-23 |
| 1.0E+00 | 1.3E-08 | 6.7E-23 | 2.6E-04 | 6.8E-03 | 6.5E-27 | 3.1E-58 |
| 1.3E-08 | 1.0E+00 | 6.0E-04 | 1.2E-01 | 1.5E-18 | 2.4E-46 | 4.5E-92 |
| 6.7E-23 | 6.0E-04 | 1.0E+00 | 2.8E-08 | 1.1E-36 | 1.0E-67 | 4.7E-128 |
| 2.6E-04 | 1.2E-01 | 2.8E-08 | 1.0E+00 | 6.2E-12 | 2.4E-46 | 4.5E-92 |
| 6.8E-03 | 1.5E-18 | 1.1E-36 | 6.2E-12 | 1.0E+00 | 4.4E-15 | 1.8E-34 |
| 6.5E-27 | 2.4E-46 | 1.0E-67 | 2.4E-46 | 4.4E-15 | 1.0E+00 | 9.9E-02 |
| 3.1E-58 | 4.5E-92 | 4.7E-128 | 4.5E-92 | 1.8E-34 | 9.9E-02 | 1.0E+00 |
| 5.5E-64 | 2.8E-99 | 1.6E-136 | 2.8E-99 | 1.2E-38 | 1.6E-02 | 1.0E+00 |
| 5.3E-64 | 1.0E-98 | 2.9E-135 | 1.0E-98 | 5.0E-39 | 8.9E-03 | 9.5E-01 |
| 5.7E-65 | 1.6E-100 | 5.7E-138 | 1.6E-100 | 2.3E-39 | 1.1E-02 | 9.8E-01 |

| 10 min | 15 min | 20 min | 25 min | 30 min | 35 min | 40 min |
|---|---|---|---|---|---|---|
| 3.6E-23 | 1.0E-44 | 2.5E-78 | 2.0E-20 | 9.1E-67 | 3.1E-21 | 1.6E-04 |
| 7.2E-26 | 3.6E-49 | 3.5E-88 | 2.7E-21 | 8.6E-74 | 1.8E-23 | 5.6E-05 |
| 5.9E-21 | 5.7E-41 | 2.3E-70 | 1.3E-19 | 6.7E-61 | 2.2E-19 | 3.8E-04 |
| 4.7E-15 | 2.0E-30 | 1.5E-49 | 7.3E-17 | 1.2E-44 | 3.0E-14 | 4.1E-03 |
| 1.9E-15 | 2.7E-35 | 4.8E-67 | 2.7E-15 | 4.3E-56 | 4.9E-14 | 2.0E-01 |
| 5.5E-05 | 4.2E-17 | 4.9E-34 | 2.0E-08 | 1.2E-29 | 1.2E-04 | 1.0E+00 |
| 1.0E+00 | 1.3E-05 | 1.1E-26 | 2.3E-03 | 2.7E-15 | 5.1E-01 | 1.3E-07 |
| 1.3E-05 | 1.0E+00 | 4.2E-06 | 1.0E+00 | 2.0E-02 | 3.4E-05 | 1.2E-22 |
| 1.1E-26 | 4.2E-06 | 1.0E+00 | 2.4E-02 | 3.3E-01 | 3.9E-17 | 6.0E-45 |
| 2.3E-03 | 1.0E+00 | 2.4E-02 | 1.0E+00 | 2.8E-01 | 1.6E-02 | 2.1E-10 |
| 2.7E-15 | 2.0E-02 | 3.3E-01 | 2.8E-01 | 1.0E+00 | 5.9E-13 | 2.5E-38 |
| 5.1E-01 | 3.4E-05 | 3.9E-17 | 1.6E-02 | 5.9E-13 | 1.0E+00 | 5.0E-07 |
| 1.3E-07 | 1.2E-22 | 6.0E-45 | 2.1E-10 | 2.5E-38 | 5.0E-07 | 1.0E+00 |
| 2.5E-14 | 2.9E-29 | 1.1E-47 | 2.4E-16 | 4.4E-43 | 1.4E-13 | 7.1E-03 |
| 5.7E-25 | 5.0E-48 | 1.9E-86 | 8.6E-21 | 2.4E-72 | 1.2E-22 | 1.3E-04 |
| 5.7E-25 | 5.0E-48 | 1.9E-86 | 8.6E-21 | 2.4E-72 | 1.2E-22 | 1.3E-04 |

| 10 min | 15 min | 20 min | 25 min | 30 min | 35 min | 40 min |
|---|---|---|---|---|---|---|
| 3.4E-165 | 7.8E-163 | 5.1E-194 | 4.6E-188 | 1.9E-147 | 4.6E-62 | 9.8E-19 |
| 1.3E-163 | 2.9E-161 | 2.6E-192 | 2.2E-186 | 4.7E-146 | 2.9E-61 | 3.3E-18 |
| 4.7E-162 | 1.0E-159 | 1.9E-191 | 1.0E-184 | 1.2E-144 | 1.8E-60 | 1.1E-17 |
| 5.0E-146 | 1.6E-147 | 1.0E-184 | 2.2E-172 | 1.7E-128 | 2.3E-51 | 4.2E-12 |
| 4.0E-101 | 4.7E-128 | 2.6E-154 | 4.5E-143 | 2.1E-60 | 5.4E-10 | 6.2E-05 |
| 1.2E-24 | 2.1E-71 | 1.0E-74 | 8.0E-72 | 1.6E-07 | 4.8E-17 | 5.9E-57 |
| 1.0E+00 | 6.2E-17 | 5.4E-15 | 3.3E-18 | 9.1E-34 | 1.2E-61 | 9.5E-129 |
| 6.2E-17 | 1.0E+00 | 1.6E-02 | 1.9E-01 | 1.5E-77 | 3.5E-80 | 7.1E-141 |
| 5.4E-15 | 1.6E-02 | 1.0E+00 | 6.1E-01 | 1.0E-84 | 1.4E-96 | 3.4E-173 |
| 3.3E-18 | 1.9E-01 | 6.1E-01 | 1.0E+00 | 7.9E-80 | 1.1E-89 | 1.3E-163 |
| 9.1E-34 | 1.5E-77 | 1.0E-84 | 7.9E-80 | 1.0E+00 | 1.9E-19 | 6.8E-95 |
| 1.2E-61 | 3.5E-80 | 1.4E-96 | 1.1E-89 | 1.9E-19 | 1.0E+00 | 5.9E-24 |
| 9.5E-129 | 7.1E-141 | 3.4E-173 | 1.3E-163 | 6.8E-95 | 5.9E-24 | 1.0E+00 |
| 5.0E-146 | 5.1E-149 | 2.2E-186 | 5.3E-174 | 1.4E-120 | 1.7E-38 | 1.8E-05 |
| 3.3E-151 | 1.0E-149 | 1.4E-186 | 3.6E-175 | 2.5E-127 | 1.5E-51 | 1.8E-12 |
| 2.8E-151 | 1.6E-150 | 4.6E-188 | 1.3E-175 | 4.1E-131 | 7.7E-53 | 5.9E-13 |

| 10 min | 15 min | 20 min | 25 min | 30 min | 35 min | 40 min |
|---|---|---|---|---|---|---|
| 2.5E-91 | 6.1E-108 | 5.1E-163 | 1.2E-58 | 2.6E-125 | 3.0E-132 | 1.8E-81 |
| 1.8E-100 | 4.2E-117 | 2.7E-181 | 2.1E-60 | 1.0E-136 | 1.7E-144 | 5.3E-89 |
| 2.3E-82 | 1.1E-98 | 3.1E-146 | 3.0E-56 | 4.0E-114 | 2.5E-120 | 7.2E-74 |
| 8.8E-56 | 6.8E-70 | 1.1E-98 | 1.7E-47 | 6.1E-81 | 2.9E-84 | 1.1E-50 |
| 3.9E-48 | 2.4E-80 | 2.5E-144 | 2.7E-45 | 6.2E-102 | 5.3E-79 | 3.9E-27 |
| 1.0E-21 | 2.7E-51 | 2.6E-96 | 6.5E-35 | 4.9E-64 | 4.2E-32 | 2.8E-03 |
| 1.0E+00 | 9.5E-25 | 1.8E-72 | 9.6E-17 | 2.9E-42 | 2.5E-06 | 4.3E-24 |
| 9.5E-25 | 1.0E+00 | 7.5E-10 | 3.3E-01 | 1.6E-04 | 2.2E-14 | 8.0E-56 |
| 1.8E-72 | 7.5E-10 | 1.0E+00 | 1.4E-02 | 3.8E-03 | 3.9E-49 | 5.7E-108 |
| 9.6E-17 | 3.3E-01 | 1.4E-02 | 1.0E+00 | 2.6E-01 | 1.7E-11 | 1.8E-35 |
| 2.9E-42 | 1.6E-04 | 3.8E-03 | 2.6E-01 | 1.0E+00 | 5.7E-29 | 9.0E-72 |
| 2.5E-06 | 2.2E-14 | 3.9E-49 | 1.7E-11 | 5.7E-29 | 1.0E+00 | 2.5E-32 |
| 4.3E-24 | 8.0E-56 | 5.7E-108 | 1.8E-35 | 9.0E-72 | 2.5E-32 | 1.0E+00 |
| 2.0E-49 | 1.2E-65 | 9.9E-93 | 1.9E-46 | 2.1E-78 | 2.2E-76 | 8.8E-45 |
| 3.0E-77 | 1.5E-102 | 7.7E-166 | 6.2E-58 | 4.1E-130 | 2.9E-115 | 1.1E-59 |
| 7.7E-89 | 5.8E-108 | 3.1E-168 | 1.7E-58 | 9.2E-132 | 3.8E-131 | 2.1E-78 |

| 10 min | 15 min | 20 min | 25 min | 30 min | 35 min | 40 min |
|---|---|---|---|---|---|---|
| 1.1E-55 | 7.8E-77 | 6.8E-49 | 1.7E-43 | 4.7E-30 | 1.3E-06 | 3.0E-09 |
| 5.8E-52 | 7.9E-72 | 1.4E-44 | 4.0E-40 | 5.7E-27 | 1.6E-05 | 1.1E-07 |
| 7.3E-48 | 2.1E-63 | 4.3E-39 | 4.5E-37 | 8.7E-25 | 7.4E-06 | 9.0E-08 |
| 9.5E-47 | 5.2E-64 | 2.0E-38 | 1.3E-35 | 4.4E-23 | 1.7E-04 | 4.2E-06 |
| 1.0E-11 | 2.0E-16 | 1.8E-04 | 2.2E-06 | 1.3E-01 | 4.0E-01 | 1.6E-01 |
| 9.9E-01 | 2.9E-02 | 1.7E-04 | 1.8E-01 | 2.4E-08 | 5.4E-16 | 2.7E-24 |
| 1.0E+00 | 5.8E-01 | 1.1E-03 | 5.7E-02 | 1.0E-06 | 1.1E-13 | 9.1E-19 |
| 5.8E-01 | 1.0E+00 | 7.0E-06 | 6.6E-05 | 7.7E-10 | 1.7E-17 | 1.7E-25 |
| 1.1E-03 | 7.0E-06 | 1.0E+00 | 5.0E-01 | 2.4E-01 | 1.1E-06 | 3.7E-10 |
| 5.7E-02 | 6.6E-05 | 5.0E-01 | 1.0E+00 | 8.0E-03 | 1.3E-08 | 5.7E-12 |
| 1.0E-06 | 7.7E-10 | 2.4E-01 | 8.0E-03 | 1.0E+00 | 3.1E-03 | 1.1E-04 |
| 1.1E-13 | 1.7E-17 | 1.1E-06 | 1.3E-08 | 3.1E-03 | 1.0E+00 | 1.0E+00 |
| 9.1E-19 | 1.7E-25 | 3.7E-10 | 5.7E-12 | 1.1E-04 | 1.0E+00 | 1.0E+00 |
| 1.1E-17 | 3.4E-25 | 6.1E-09 | 1.0E-10 | 1.6E-03 | 1.0E+00 | 9.5E-01 |
| 2.1E-14 | 9.3E-19 | 4.8E-07 | 5.5E-09 | 2.7E-03 | 1.0E+00 | 1.0E+00 |
| 6.7E-09 | 2.5E-13 | 2.7E-02 | 4.5E-04 | 6.6E-01 | 9.6E-03 | 3.9E-04 |

| 10 min | 15 min | 20 min | 25 min | 30 min | 35 min | 40 min |
|---|---|---|---|---|---|---|
| 5.3E-24 | 4.7E-53 | 3.5E-34 | 1.3E-12 | 3.3E-04 | 7.1E-05 | 3.0E-03 |
| 5.3E-24 | 4.7E-53 | 3.5E-34 | 1.3E-12 | 3.3E-04 | 7.1E-05 | 3.0E-03 |
| 5.3E-24 | 4.7E-53 | 3.5E-34 | 1.3E-12 | 3.3E-04 | 7.1E-05 | 3.0E-03 |
| 1.8E-11 | 3.5E-33 | 1.0E-18 | 1.6E-06 | 4.1E-01 | 8.4E-01 | 1.0E+00 |
| 1.0E+00 | 3.5E-06 | 2.5E-01 | 7.1E-01 | 6.1E-03 | 5.8E-08 | 1.3E-09 |
| 2.6E-12 | 1.3E-01 | 1.0E-06 | 1.1E-04 | 3.1E-17 | 2.8E-38 | 1.3E-40 |
| 1.0E+00 | 3.5E-06 | 2.5E-01 | 6.4E-01 | 6.1E-03 | 5.8E-08 | 1.3E-09 |
| 3.5E-06 | 1.0E+00 | 1.6E-02 | 3.1E-02 | 1.6E-11 | 6.8E-27 | 3.6E-29 |
| 2.5E-01 | 1.6E-02 | 1.0E+00 | 1.0E+00 | 1.3E-05 | 4.1E-14 | 4.3E-16 |
| 6.4E-01 | 3.1E-02 | 1.0E+00 | 1.0E+00 | 1.6E-02 | 9.0E-05 | 9.6E-06 |
| 6.1E-03 | 1.6E-11 | 1.3E-05 | 1.6E-02 | 1.0E+00 | 9.9E-01 | 6.7E-01 |
| 5.8E-08 | 6.8E-27 | 4.1E-14 | 9.0E-05 | 9.9E-01 | 1.0E+00 | 1.0E+00 |
| 1.3E-09 | 3.6E-29 | 4.3E-16 | 9.6E-06 | 6.7E-01 | 1.0E+00 | 1.0E+00 |
| 4.4E-07 | 1.2E-22 | 3.5E-12 | 1.0E-04 | 9.5E-01 | 1.0E+00 | 1.0E+00 |
| 7.7E-03 | 2.2E-08 | 1.2E-04 | 1.0E-02 | 1.0E+00 | 1.0E+00 | 1.0E+00 |
| 5.3E-05 | 5.1E-21 | 5.4E-10 | 3.0E-03 | 1.0E+00 | 7.9E-01 | 2.4E-01 |

| 10 min | 15 min | 20 min | 25 min | 30 min | 35 min | 40 min |
|---|---|---|---|---|---|---|
| 2.1E-153 | 1.9E-233 | 2.1E-290 | 9.6E-195 | 2.5E-249 | 2.8E-98 | 4.1E-110 |
| 8.9E-151 | 8.1E-234 | 2.1E-290 | 1.1E-192 | 2.3E-240 | 2.7E-93 | 1.1E-103 |
| 6.8E-132 | 3.7E-196 | 4.3E-236 | 5.7E-167 | 1.2E-204 | 2.0E-86 | 1.4E-91 |
| 6.4E-128 | 3.5E-196 | 3.5E-242 | 2.6E-163 | 4.7E-202 | 7.5E-71 | 2.2E-74 |
| 1.3E-61 | 9.2E-95 | 2.2E-116 | 4.8E-81 | 1.5E-100 | 2.3E-34 | 1.0E-38 |
| 5.4E-02 | 1.1E-01 | 2.6E-03 | 9.7E-04 | 1.3E-08 | 2.2E-28 | 2.9E-38 |
| 1.0E+00 | 5.1E-02 | 1.1E-02 | 3.9E-06 | 3.7E-11 | 1.3E-26 | 5.9E-35 |
| 5.1E-02 | 1.0E+00 | 1.2E-01 | 2.7E-04 | 9.6E-09 | 2.1E-32 | 3.8E-40 |
| 1.1E-02 | 1.2E-01 | 1.0E+00 | 5.8E-07 | 7.8E-14 | 2.0E-41 | 1.6E-53 |
| 3.9E-06 | 2.7E-04 | 5.8E-07 | 1.0E+00 | 2.7E-01 | 1.1E-19 | 3.0E-29 |
| 3.7E-11 | 9.6E-09 | 7.8E-14 | 2.7E-01 | 1.0E+00 | 2.6E-20 | 5.7E-34 |
| 1.3E-26 | 2.1E-32 | 2.0E-41 | 1.1E-19 | 2.6E-20 | 1.0E+00 | 2.5E-02 |
| 5.9E-35 | 3.8E-40 | 1.6E-53 | 3.0E-29 | 5.7E-34 | 2.5E-02 | 1.0E+00 |
| 2.2E-39 | 7.0E-52 | 1.2E-69 | 1.3E-34 | 1.9E-41 | 5.4E-02 | 7.6E-01 |
| 8.2E-28 | 5.8E-40 | 4.0E-49 | 6.2E-32 | 6.6E-36 | 5.3E-06 | 1.9E-02 |
| 1.4E-27 | 1.3E-35 | 5.3E-50 | 1.7E-24 | 1.7E-29 | 2.0E-02 | 3.3E-02 |

| 10 min | 15 min | 20 min | 25 min | 30 min | 35 min | 40 min |
|---|---|---|---|---|---|---|
| 1.2E-116 | 1.5E-141 | 6.7E-163 | 1.0E-46 | 2.2E-56 | 5.7E-86 | 1.2E-55 |
| 3.0E-124 | 3.4E-149 | 4.2E-171 | 1.8E-50 | 7.3E-61 | 1.3E-93 | 7.7E-64 |
| 5.7E-123 | 2.2E-146 | 4.3E-168 | 6.3E-50 | 3.3E-60 | 1.7E-92 | 3.6E-58 |
| 6.2E-104 | 1.7E-127 | 8.6E-148 | 3.4E-41 | 2.2E-49 | 4.1E-74 | 3.2E-40 |
| 8.6E-28 | 1.0E-42 | 6.7E-54 | 5.5E-10 | 8.1E-10 | 2.7E-09 | 6.7E-04 |
| 5.6E-01 | 7.1E-07 | 6.2E-10 | 5.5E-02 | 5.9E-04 | 1.7E-21 | 4.3E-38 |
| 1.0E+00 | 2.4E-07 | 4.7E-12 | 7.9E-02 | 1.1E-03 | 4.9E-17 | 1.8E-32 |
| 2.4E-07 | 1.0E+00 | 2.5E-01 | 1.5E-06 | 3.4E-12 | 1.7E-40 | 1.0E-59 |
| 4.7E-12 | 2.5E-01 | 1.0E+00 | 7.1E-10 | 6.5E-16 | 1.2E-52 | 9.3E-73 |
| 7.9E-02 | 1.5E-06 | 7.1E-10 | 1.0E+00 | 4.1E-01 | 6.9E-04 | 1.8E-09 |
| 1.1E-03 | 3.4E-12 | 6.5E-16 | 4.1E-01 | 1.0E+00 | 3.1E-03 | 1.8E-11 |
| 4.9E-17 | 1.7E-40 | 1.2E-52 | 6.9E-04 | 3.1E-03 | 1.0E+00 | 2.0E-08 |
| 1.8E-32 | 1.0E-59 | 9.3E-73 | 1.8E-09 | 1.8E-11 | 2.0E-08 | 1.0E+00 |
| 1.1E-28 | 7.1E-48 | 6.8E-58 | 2.9E-10 | 9.8E-12 | 9.4E-09 | 3.0E-01 |
| 5.8E-13 | 4.5E-20 | 4.2E-24 | 1.0E-07 | 1.3E-07 | 3.5E-05 | 1.6E-01 |
| 1.0E-18 | 1.7E-40 | 1.2E-52 | 1.5E-05 | 4.2E-05 | 2.0E-03 | 1.8E-03 |

| 10 min | 15 min | 20 min | 25 min | 30 min | 35 min | 40 min |
|---|---|---|---|---|---|---|
| 5.6E-68 | 1.2E-92 | 1.6E-150 | 1.9E-94 | 7.5E-145 | 3.6E-47 | 4.0E-47 |
| 1.5E-71 | 7.5E-97 | 7.5E-156 | 1.1E-98 | 9.4E-150 | 1.2E-49 | 4.1E-50 |
| 1.8E-68 | 2.9E-93 | 2.8E-151 | 4.8E-95 | 1.5E-145 | 1.6E-47 | 1.5E-47 |
| 1.3E-60 | 5.2E-84 | 2.0E-139 | 1.0E-85 | 1.3E-134 | 4.3E-42 | 5.0E-41 |
| 1.9E-30 | 1.5E-47 | 6.7E-91 | 8.0E-49 | 1.2E-89 | 2.6E-21 | 3.5E-17 |
| 2.2E-06 | 1.6E-14 | 4.8E-49 | 6.8E-15 | 3.0E-61 | 2.6E-04 | 9.8E-03 |
| 1.0E+00 | 2.4E-02 | 3.5E-21 | 1.3E-02 | 4.9E-31 | 7.3E-01 | 2.4E-02 |
| 2.4E-02 | 1.0E+00 | 2.6E-12 | 5.0E-01 | 2.1E-21 | 1.9E-03 | 4.2E-08 |
| 3.5E-21 | 2.6E-12 | 1.0E+00 | 1.9E-16 | 3.1E-02 | 6.7E-20 | 1.9E-30 |
| 1.3E-02 | 5.0E-01 | 1.9E-16 | 1.0E+00 | 7.4E-26 | 7.9E-02 | 1.3E-08 |
| 4.9E-31 | 2.1E-21 | 3.1E-02 | 7.4E-26 | 1.0E+00 | 2.5E-28 | 2.3E-41 |
| 7.3E-01 | 1.9E-03 | 6.7E-20 | 7.9E-02 | 2.5E-28 | 1.0E+00 | 7.9E-02 |
| 2.4E-02 | 4.2E-08 | 1.9E-30 | 1.3E-08 | 2.3E-41 | 7.9E-02 | 1.0E+00 |
| 8.6E-03 | 5.6E-09 | 1.8E-31 | 1.6E-09 | 9.6E-42 | 3.8E-02 | 1.0E+00 |
| 2.7E-12 | 1.4E-23 | 1.8E-55 | 1.9E-24 | 1.7E-56 | 7.5E-09 | 9.3E-05 |
| 2.7E-14 | 1.5E-26 | 1.3E-60 | 1.7E-27 | 1.3E-65 | 3.7E-10 | 6.7E-06 |

| 10 min | 15 min | 20 min | 25 min | 30 min | 35 min | 40 min |
|---|---|---|---|---|---|---|
| 4.0E-46 | 5.5E-74 | 1.7E-97 | 3.2E-42 | 8.8E-73 | 4.0E-88 | 2.4E-75 |
| 6.0E-55 | 2.7E-85 | 7.4E-115 | 6.4E-46 | 1.9E-84 | 6.1E-102 | 7.0E-88 |
| 3.1E-44 | 8.7E-71 | 2.3E-91 | 3.7E-42 | 1.9E-69 | 1.1E-83 | 1.3E-71 |
| 3.5E-28 | 3.5E-48 | 3.5E-59 | 1.7E-33 | 1.3E-46 | 2.1E-56 | 1.8E-47 |
| 1.1E-22 | 4.6E-46 | 1.5E-63 | 1.2E-26 | 1.4E-44 | 2.8E-57 | 3.4E-46 |
| 8.4E-03 | 2.4E-13 | 9.8E-19 | 3.6E-13 | 9.7E-14 | 8.7E-27 | 6.3E-21 |
| 1.0E+00 | 1.3E-08 | 2.7E-13 | 1.9E-09 | 6.3E-09 | 4.0E-21 | 1.8E-15 |
| 1.3E-08 | 1.0E+00 | 1.0E+00 | 1.6E-01 | 8.6E-01 | 1.2E-02 | 4.3E-01 |
| 2.7E-13 | 1.0E+00 | 1.0E+00 | 3.7E-01 | 7.5E-01 | 5.1E-02 | 6.2E-01 |
| 1.9E-09 | 1.6E-01 | 3.7E-01 | 1.0E+00 | 1.4E-01 | 5.0E-01 | 3.7E-01 |
| 6.3E-09 | 8.6E-01 | 7.5E-01 | 1.4E-01 | 1.0E+00 | 4.4E-03 | 1.2E-01 |
| 4.0E-21 | 1.2E-02 | 5.1E-02 | 5.0E-01 | 4.4E-03 | 1.0E+00 | 3.6E-01 |
| 1.8E-15 | 4.3E-01 | 6.2E-01 | 3.7E-01 | 1.2E-01 | 3.6E-01 | 1.0E+00 |
| 8.4E-01 | 1.5E-07 | 2.2E-10 | 2.8E-09 | 1.1E-07 | 2.8E-16 | 3.2E-12 |
| 6.9E-03 | 2.6E-15 | 1.6E-22 | 1.1E-10 | 6.6E-14 | 1.7E-24 | 2.7E-18 |
| 2.5E-14 | 1.4E-34 | 2.2E-48 | 7.7E-21 | 4.9E-33 | 5.6E-44 | 3.9E-34 |

| 10 min | 15 min | 20 min | 25 min | 30 min | 35 min | 40 min |
|---|---|---|---|---|---|---|
| 1.6E-150 | 9.3E-190 | 3.4E-199 | 1.3E-190 | 3.9E-168 | 5.1E-128 | 1.3E-163 |
| 1.6E-150 | 1.9E-191 | 4.7E-200 | 1.9E-191 | 6.9E-169 | 1.4E-128 | 1.3E-163 |
| 1.6E-150 | 6.5E-189 | 1.9E-197 | 6.5E-189 | 8.4E-168 | 7.2E-127 | 2.1E-164 |
| 1.5E-144 | 3.2E-187 | 1.9E-197 | 2.2E-186 | 2.0E-164 | 3.6E-124 | 2.2E-157 |
| 7.5E-94 | 8.6E-153 | 3.6E-169 | 3.4E-173 | 1.3E-129 | 1.2E-101 | 4.9E-108 |
| 8.0E-32 | 3.3E-87 | 2.9E-101 | 4.9E-125 | 1.2E-59 | 5.0E-59 | 5.8E-43 |
| 1.0E+00 | 2.6E-22 | 1.1E-24 | 9.4E-53 | 6.1E-09 | 7.5E-15 | 4.6E-04 |
| 2.6E-22 | 1.0E+00 | 4.5E-01 | 1.1E-11 | 8.9E-04 | 7.9E-01 | 1.6E-09 |
| 1.1E-24 | 4.5E-01 | 1.0E+00 | 2.6E-12 | 1.8E-04 | 2.3E-01 | 1.3E-13 |
| 9.4E-53 | 1.1E-11 | 2.6E-12 | 1.0E+00 | 4.4E-20 | 2.5E-06 | 1.7E-32 |
| 6.1E-09 | 8.9E-04 | 1.8E-04 | 4.4E-20 | 1.0E+00 | 4.0E-03 | 1.1E-03 |
| 7.5E-15 | 7.9E-01 | 2.3E-01 | 2.5E-06 | 4.0E-03 | 1.0E+00 | 3.5E-06 |
| 4.6E-04 | 1.6E-09 | 1.3E-13 | 1.7E-32 | 1.1E-03 | 3.5E-06 | 1.0E+00 |
| 1.8E-05 | 5.4E-35 | 5.2E-43 | 5.7E-65 | 1.9E-21 | 3.8E-23 | 6.7E-10 |
| 6.6E-25 | 1.1E-67 | 1.8E-77 | 2.6E-103 | 3.1E-49 | 5.7E-46 | 3.2E-31 |
| 2.0E-41 | 6.7E-91 | 2.3E-102 | 7.7E-120 | 4.9E-70 | 7.2E-63 | 1.1E-49 |

| 10 min | 15 min | 20 min | 25 min | 30 min | 35 min | 40 min |
|---|---|---|---|---|---|---|
| 2.1E-92 | 8.9E-123 | 3.7E-174 | 2.5E-58 | 7.1E-141 | 5.3E-144 | 2.8E-136 |
| 2.2E-103 | 2.9E-135 | 7.6E-195 | 2.1E-60 | 9.0E-157 | 1.3E-159 | 1.6E-152 |
| 1.3E-86 | 5.7E-112 | 5.7E-154 | 4.6E-56 | 1.2E-127 | 1.1E-130 | 1.5E-122 |
| 2.2E-68 | 4.4E-87 | 3.6E-111 | 1.1E-50 | 4.4E-97 | 1.2E-99 | 3.2E-93 |
| 1.3E-60 | 7.5E-100 | 3.7E-177 | 1.6E-53 | 3.3E-140 | 2.4E-132 | 1.5E-114 |
| 5.1E-16 | 1.1E-55 | 2.5E-109 | 3.6E-39 | 3.6E-88 | 9.0E-80 | 4.6E-68 |
| 1.0E+00 | 1.6E-27 | 8.2E-82 | 5.8E-30 | 2.7E-68 | 1.3E-53 | 2.4E-43 |
| 1.6E-27 | 1.0E+00 | 2.8E-15 | 1.5E-07 | 6.2E-21 | 3.4E-13 | 6.9E-08 |
| 8.2E-82 | 2.8E-15 | 1.0E+00 | 7.1E-01 | 8.8E-04 | 7.7E-03 | 2.4E-06 |
| 5.8E-30 | 1.5E-07 | 7.1E-01 | 1.0E+00 | 8.4E-02 | 4.1E-01 | 2.9E-02 |
| 2.7E-68 | 6.2E-21 | 8.8E-04 | 8.4E-02 | 1.0E+00 | 2.7E-02 | 4.8E-06 |
| 1.3E-53 | 3.4E-13 | 7.7E-03 | 4.1E-01 | 2.7E-02 | 1.0E+00 | 1.6E-02 |
| 2.4E-43 | 6.9E-08 | 2.4E-06 | 2.9E-02 | 4.8E-06 | 1.6E-02 | 1.0E+00 |
| 3.9E-03 | 5.2E-09 | 5.0E-32 | 5.4E-17 | 2.9E-34 | 2.8E-24 | 7.0E-17 |
| 4.8E-16 | 7.9E-09 | 2.7E-35 | 1.8E-10 | 2.4E-27 | 8.6E-19 | 2.1E-12 |
| 4.7E-05 | 5.4E-35 | 1.9E-86 | 5.1E-25 | 3.1E-64 | 2.6E-56 | 3.3E-45 |

| 45 min | 50 min | 55 min |
|---|---|---|
| 1.0E+00 | 1.0E+00 | 1.0E+00 |
| 1.0E+00 | 1.0E+00 | 1.0E+00 |
| 1.0E+00 | 1.0E+00 | 1.0E+00 |
| 1.0E+00 | 1.0E+00 | 1.0E+00 |
| 4.7E-11 | 2.0E-07 | 2.6E-11 |
| 1.3E-76 | 8.1E-47 | 2.4E-80 |
| 3.5E-71 | 4.8E-50 | 2.5E-73 |
| 3.0E-99 | 2.7E-64 | 2.2E-103 |
| 8.2E-30 | 2.5E-18 | 4.3E-31 |
| 2.1E-02 | 8.6E-02 | 2.1E-02 |
| 1.0E+00 | 1.0E+00 | 1.0E+00 |
| 1.0E+00 | 1.0E+00 | 1.0E+00 |
| 1.0E+00 | 1.0E+00 | 1.0E+00 |
| 1.0E+00 | 1.0E+00 | 1.0E+00 |
| 1.0E+00 | 1.0E+00 | 1.0E+00 |
| 1.0E+00 | 1.0E+00 | 1.0E+00 |

| 45 min | 50 min | 55 min |
|---|---|---|
| 1.0E+00 | 1.0E+00 | 1.0E+00 |
| 1.0E+00 | 1.0E+00 | 1.0E+00 |
| 1.0E+00 | 1.0E+00 | 1.0E+00 |
| 8.6E-01 | 1.0E+00 | 7.4E-01 |
| 2.0E-07 | 6.8E-03 | 2.7E-09 |
| 3.3E-22 | 3.2E-08 | 8.5E-28 |
| 4.1E-27 | 7.8E-10 | 3.5E-34 |
| 2.1E-40 | 1.6E-14 | 5.6E-51 |
| 5.2E-09 | 1.9E-03 | 2.8E-11 |
| 2.3E-01 | 6.2E-01 | 1.8E-01 |
| 1.0E+00 | 1.0E+00 | 1.0E+00 |
| 1.0E+00 | 1.0E+00 | 1.0E+00 |
| 1.0E+00 | 1.0E+00 | 1.0E+00 |
| 1.0E+00 | 1.0E+00 | 1.0E+00 |
| 1.0E+00 | 1.0E+00 | 1.0E+00 |

| | | |
|---|---|---|
| 1.0E+00 | 1.0E+00 | 1.0E+00 |

| 45 min | 50 min | 55 min |
|---|---|---|
| 1.0E+00 | 1.0E+00 | 1.0E+00 |
| 9.9E-01 | 1.0E+00 | 1.0E+00 |
| 9.9E-01 | 1.0E+00 | 1.0E+00 |
| 9.9E-01 | 1.0E+00 | 1.0E+00 |
| 6.9E-38 | 4.4E-25 | 4.2E-41 |
| 9.4E-241 | 1.1E-146 | 5.7E-259 |
| 4.7E-148 | 7.0E-104 | 2.8E-156 |
| 8.0E-259 | 2.6E-167 | 1.5E-275 |
| 5.6E-229 | 1.6E-139 | 2.1E-246 |
| 6.8E-48 | 4.1E-33 | 2.4E-51 |
| 6.0E-08 | 2.9E-05 | 5.3E-09 |
| 1.0E+00 | 1.0E+00 | 1.0E+00 |
| 1.0E+00 | 1.0E+00 | 1.0E+00 |
| 1.0E+00 | 1.0E+00 | 1.0E+00 |
| 1.0E+00 | 1.0E+00 | 1.0E+00 |
| 1.0E+00 | 1.0E+00 | 1.0E+00 |

| 45 min | 50 min | 55 min |
|---|---|---|
| 1.0E+00 | 1.0E+00 | 1.0E+00 |
| 1.0E+00 | 1.0E+00 | 1.0E+00 |
| 1.0E+00 | 1.0E+00 | 1.0E+00 |
| 1.1E-01 | 6.8E-01 | 1.2E-01 |
| 1.3E-20 | 1.6E-07 | 2.8E-24 |
| 1.3E-59 | 1.3E-21 | 5.6E-72 |
| 7.2E-88 | 3.1E-31 | 1.7E-107 |
| 6.9E-103 | 1.6E-36 | 3.4E-126 |
| 9.3E-92 | 1.3E-32 | 2.5E-112 |
| 5.7E-11 | 6.3E-06 | 2.0E-11 |
| 1.8E-02 | 2.1E-01 | 2.2E-02 |
| 5.5E-01 | 9.8E-01 | 6.9E-01 |
| 1.0E+00 | 1.0E+00 | 1.0E+00 |
| 1.0E+00 | 1.0E+00 | 1.0E+00 |
| 1.0E+00 | 1.0E+00 | 1.0E+00 |
| 1.0E+00 | 1.0E+00 | 1.0E+00 |

| 45 min | 50 min | 55 min |
|---|---|---|
| 1.0E+00 | 1.0E+00 | 1.0E+00 |
| 1.0E+00 | 1.0E+00 | 1.0E+00 |
| 1.0E+00 | 1.0E+00 | 1.0E+00 |
| 1.0E+00 | 1.0E+00 | 1.0E+00 |
| 4.3E-02 | 2.3E-02 | 2.9E-02 |
| 2.2E-26 | 7.9E-27 | 5.2E-27 |
| 5.5E-64 | 5.3E-64 | 5.7E-65 |
| 2.8E-99 | 1.0E-98 | 1.6E-100 |
| 1.6E-136 | 2.9E-135 | 5.7E-138 |
| 2.8E-99 | 1.0E-98 | 1.6E-100 |
| 1.2E-38 | 5.0E-39 | 2.3E-39 |
| 1.6E-02 | 8.9E-03 | 1.1E-02 |
| 1.0E+00 | 9.5E-01 | 9.8E-01 |
| 1.0E+00 | 1.0E+00 | 1.0E+00 |
| 1.0E+00 | 1.0E+00 | 1.0E+00 |
| 1.0E+00 | 1.0E+00 | 1.0E+00 |

| 45 min | 50 min | 55 min |
|---|---|---|
| 1.0E+00 | 1.0E+00 | 1.0E+00 |
| 1.0E+00 | 1.0E+00 | 1.0E+00 |
| 1.0E+00 | 1.0E+00 | 1.0E+00 |
| 1.0E+00 | 1.0E+00 | 1.0E+00 |
| 4.0E-01 | 1.2E-01 | 1.2E-01 |
| 1.4E-03 | 1.8E-05 | 1.8E-05 |
| 2.5E-14 | 5.7E-25 | 5.7E-25 |
| 2.9E-29 | 5.0E-48 | 5.0E-48 |
| 1.1E-47 | 1.9E-86 | 1.9E-86 |
| 2.4E-16 | 8.6E-21 | 8.6E-21 |
| 4.4E-43 | 2.4E-72 | 2.4E-72 |
| 1.4E-13 | 1.2E-22 | 1.2E-22 |
| 7.1E-03 | 1.3E-04 | 1.3E-04 |
| 1.0E+00 | 1.0E+00 | 1.0E+00 |
| 1.0E+00 | 1.0E+00 | 1.0E+00 |
| 1.0E+00 | 1.0E+00 | 1.0E+00 |

| 45 min | 50 min | 55 min |
| --- | --- | --- |
| 1.5E-04 | 5.4E-01 | 5.6E-01 |
| 2.6E-04 | 6.5E-01 | 6.7E-01 |
| 4.6E-04 | 7.6E-01 | 7.8E-01 |
| 8.6E-02 | 9.9E-01 | 1.0E+00 |
| 1.0E-15 | 1.7E-26 | 1.7E-27 |
| 1.3E-77 | 8.0E-91 | 6.7E-94 |
| 5.0E-146 | 3.3E-151 | 2.8E-151 |
| 5.1E-149 | 1.0E-149 | 1.6E-150 |
| 2.2E-186 | 1.4E-186 | 4.6E-188 |
| 5.3E-174 | 3.6E-175 | 1.3E-175 |
| 1.4E-120 | 2.5E-127 | 4.1E-131 |
| 1.7E-38 | 1.5E-51 | 7.7E-53 |
| 1.8E-05 | 1.8E-12 | 5.9E-13 |
| 1.0E+00 | 5.3E-02 | 4.3E-02 |
| 5.3E-02 | 1.0E+00 | 1.0E+00 |
| 4.3E-02 | 1.0E+00 | 1.0E+00 |

| 45 min | 50 min | 55 min |
| --- | --- | --- |
| 8.6E-01 | 1.2E-03 | 6.9E-01 |
| 9.1E-01 | 9.4E-04 | 7.7E-01 |
| 8.9E-01 | 2.6E-03 | 7.7E-01 |
| 1.0E+00 | 6.6E-02 | 1.0E+00 |
| 2.2E-08 | 1.2E-07 | 1.3E-15 |
| 5.3E-25 | 7.7E-29 | 4.5E-41 |
| 2.0E-49 | 3.0E-77 | 7.7E-89 |
| 1.2E-65 | 1.5E-102 | 5.8E-108 |
| 9.9E-93 | 7.7E-166 | 3.1E-168 |
| 1.9E-46 | 6.2E-58 | 1.7E-58 |
| 2.1E-78 | 4.1E-130 | 9.2E-132 |
| 2.2E-76 | 2.9E-115 | 3.8E-131 |
| 8.8E-45 | 1.1E-59 | 2.1E-78 |
| 1.0E+00 | 4.0E-01 | 1.0E+00 |
| 4.0E-01 | 1.0E+00 | 7.0E-02 |
| 1.0E+00 | 7.0E-02 | 1.0E+00 |

| 45 min | 50 min | 55 min |
|---|---|---|
| 3.1E-15 | 9.2E-08 | 9.4E-32 |
| 5.8E-13 | 1.6E-06 | 2.6E-28 |
| 2.5E-12 | 8.7E-07 | 2.2E-25 |
| 1.9E-10 | 2.6E-05 | 7.4E-24 |
| 7.1E-01 | 4.4E-01 | 4.7E-01 |
| 3.2E-24 | 3.0E-17 | 7.3E-12 |
| 1.1E-17 | 2.1E-14 | 6.7E-09 |
| 3.4E-25 | 9.3E-19 | 2.5E-13 |
| 6.1E-09 | 4.8E-07 | 2.7E-02 |
| 1.0E-10 | 5.5E-09 | 4.5E-04 |
| 1.6E-03 | 2.7E-03 | 6.6E-01 |
| 1.0E+00 | 1.0E+00 | 9.6E-03 |
| 9.5E-01 | 1.0E+00 | 3.9E-04 |
| 1.0E+00 | 9.4E-01 | 5.9E-03 |
| 9.4E-01 | 1.0E+00 | 8.7E-03 |
| 5.9E-03 | 8.7E-03 | 1.0E+00 |

| 45 min | 50 min | 55 min |
|---|---|---|
| 1.9E-03 | 1.9E-01 | 8.3E-08 |
| 1.9E-03 | 1.9E-01 | 8.3E-08 |
| 1.9E-03 | 1.9E-01 | 8.3E-08 |
| 1.0E+00 | 1.0E+00 | 7.9E-02 |
| 4.4E-07 | 7.7E-03 | 5.3E-05 |
| 4.5E-32 | 6.5E-12 | 2.7E-31 |
| 4.4E-07 | 7.7E-03 | 5.3E-05 |
| 1.2E-22 | 2.2E-08 | 5.1E-21 |
| 3.5E-12 | 1.2E-04 | 5.4E-10 |
| 1.0E-04 | 1.0E-02 | 3.0E-03 |
| 9.5E-01 | 1.0E+00 | 1.0E+00 |
| 1.0E+00 | 1.0E+00 | 7.9E-01 |
| 1.0E+00 | 1.0E+00 | 2.4E-01 |
| 1.0E+00 | 1.0E+00 | 6.6E-01 |
| 1.0E+00 | 1.0E+00 | 9.9E-01 |
| 6.6E-01 | 9.9E-01 | 1.0E+00 |

| 45 min | 50 min | 55 min |
|---|---|---|
| 1.5E-137 | 5.4E-58 | 3.8E-162 |
| 1.4E-129 | 2.7E-53 | 3.2E-153 |
| 2.7E-110 | 1.6E-49 | 1.2E-128 |
| 4.1E-92 | 4.7E-37 | 2.4E-110 |
| 5.5E-42 | 1.2E-17 | 1.1E-57 |
| 3.4E-47 | 8.9E-33 | 8.9E-34 |
| 2.2E-39 | 8.2E-28 | 1.4E-27 |
| 7.0E-52 | 5.8E-40 | 1.3E-35 |
| 1.2E-69 | 4.0E-49 | 5.3E-50 |
| 1.3E-34 | 6.2E-32 | 1.7E-24 |
| 1.9E-41 | 6.6E-36 | 1.7E-29 |
| 5.4E-02 | 5.3E-06 | 2.0E-02 |
| 7.6E-01 | 1.9E-02 | 3.3E-02 |
| 1.0E+00 | 6.8E-03 | 6.8E-03 |
| 6.8E-03 | 1.0E+00 | 7.4E-05 |
| 6.8E-03 | 7.4E-05 | 1.0E+00 |

| 45 min | 50 min | 55 min |
|---|---|---|
| 1.5E-41 | 3.9E-11 | 2.5E-57 |
| 4.8E-48 | 4.1E-13 | 6.4E-66 |
| 1.5E-43 | 9.7E-12 | 3.1E-62 |
| 2.0E-29 | 3.8E-08 | 2.4E-47 |
| 1.4E-02 | 1.3E-01 | 5.7E-02 |
| 6.9E-30 | 4.6E-13 | 4.5E-22 |
| 1.1E-28 | 5.8E-13 | 1.0E-18 |
| 7.1E-48 | 4.5E-20 | 1.7E-40 |
| 6.8E-58 | 4.2E-24 | 1.2E-52 |
| 2.9E-10 | 1.0E-07 | 1.5E-05 |
| 9.8E-12 | 1.3E-07 | 4.2E-05 |
| 9.4E-09 | 3.5E-05 | 2.0E-03 |
| 3.0E-01 | 1.6E-01 | 1.8E-03 |
| 1.0E+00 | 9.2E-01 | 1.8E-03 |
| 9.2E-01 | 1.0E+00 | 2.0E-02 |
| 1.8E-03 | 2.0E-02 | 1.0E+00 |

| 45 min | 50 min | 55 min |
|---|---|---|
| 4.8E-45 | 4.8E-22 | 1.3E-20 |
| 5.7E-48 | 4.6E-24 | 1.3E-22 |
| 1.9E-45 | 2.5E-22 | 6.7E-21 |
| 4.3E-39 | 5.0E-18 | 1.1E-16 |
| 6.0E-16 | 1.9E-04 | 1.7E-03 |
| 1.2E-02 | 1.0E-01 | 2.6E-02 |
| 8.6E-03 | 2.7E-12 | 2.7E-14 |
| 5.6E-09 | 1.4E-23 | 1.5E-26 |
| 1.8E-31 | 1.8E-55 | 1.3E-60 |
| 1.6E-09 | 1.9E-24 | 1.7E-27 |
| 9.6E-42 | 1.7E-56 | 1.3E-65 |
| 3.8E-02 | 7.5E-09 | 3.7E-10 |
| 1.0E+00 | 9.3E-05 | 6.7E-06 |
| 1.0E+00 | 3.8E-04 | 3.3E-05 |
| 3.8E-04 | 1.0E+00 | 5.2E-01 |
| 3.3E-05 | 5.2E-01 | 1.0E+00 |

| 45 min | 50 min | 55 min |
|---|---|---|
| 1.2E-22 | 7.6E-30 | 1.7E-11 |
| 6.1E-26 | 2.1E-36 | 1.2E-14 |
| 9.7E-23 | 6.0E-29 | 1.1E-11 |
| 5.1E-16 | 1.2E-17 | 1.3E-06 |
| 1.6E-09 | 5.8E-11 | 1.8E-01 |
| 6.3E-01 | 3.8E-01 | 1.7E-03 |
| 8.4E-01 | 6.9E-03 | 2.5E-14 |
| 1.5E-07 | 2.6E-15 | 1.4E-34 |
| 2.2E-10 | 1.6E-22 | 2.2E-48 |
| 2.8E-09 | 1.1E-10 | 7.7E-21 |
| 1.1E-07 | 6.6E-14 | 4.9E-33 |
| 2.8E-16 | 1.7E-24 | 5.6E-44 |
| 3.2E-12 | 2.7E-18 | 3.9E-34 |
| 1.0E+00 | 6.8E-01 | 1.6E-05 |
| 6.8E-01 | 1.0E+00 | 2.2E-05 |
| 1.6E-05 | 2.2E-05 | 1.0E+00 |

| 45 min | 50 min | 55 min |
|---|---|---|
| 4.2E-135 | 4.6E-96 | 1.9E-83 |
| 4.2E-135 | 4.2E-103 | 1.2E-92 |
| 4.2E-135 | 1.7E-102 | 4.5E-92 |
| 1.9E-129 | 4.7E-100 | 9.9E-90 |
| 1.6E-62 | 4.2E-31 | 2.9E-19 |
| 9.8E-14 | 3.2E-02 | 7.3E-04 |
| 1.8E-05 | 6.6E-25 | 2.0E-41 |
| 5.4E-35 | 1.1E-67 | 6.7E-91 |
| 5.2E-43 | 1.8E-77 | 2.3E-102 |
| 5.7E-65 | 2.6E-103 | 7.7E-120 |
| 1.9E-21 | 3.1E-49 | 4.9E-70 |
| 3.8E-23 | 5.7E-46 | 7.2E-63 |
| 6.7E-10 | 3.2E-31 | 1.1E-49 |
| 1.0E+00 | 1.4E-08 | 4.5E-20 |
| 1.4E-08 | 1.0E+00 | 1.8E-03 |
| 4.5E-20 | 1.8E-03 | 1.0E+00 |

| 45 min | 50 min | 55 min |
|---|---|---|
| 1.1E-82 | 2.5E-119 | 1.4E-80 |
| 2.3E-88 | 4.0E-134 | 3.6E-84 |
| 4.5E-79 | 3.3E-111 | 1.4E-71 |
| 1.6E-67 | 7.9E-88 | 3.0E-61 |
| 2.3E-57 | 6.3E-82 | 3.4E-44 |
| 2.9E-17 | 4.0E-30 | 1.1E-06 |
| 3.9E-03 | 4.8E-16 | 4.7E-05 |
| 5.2E-09 | 7.9E-09 | 5.4E-35 |
| 5.0E-32 | 2.7E-35 | 1.9E-86 |
| 5.4E-17 | 1.8E-10 | 5.1E-25 |
| 2.9E-34 | 2.4E-27 | 3.1E-64 |
| 2.8E-24 | 8.6E-19 | 2.6E-56 |
| 7.0E-17 | 2.1E-12 | 3.3E-45 |
| 1.0E+00 | 5.5E-05 | 2.0E-08 |
| 5.5E-05 | 1.0E+00 | 1.2E-12 |
| 2.0E-08 | 1.2E-12 | 1.0E+00 |
